# Human organoid screen links transient translational modulation to progenitor progression in corticogenesis

**DOI:** 10.64898/2026.11.05.764664

**Authors:** Merve Avar, Travis C. Lantz, Daniel Heinzer, Massimo M. Onesto, Kent Imaizumi, Vrishika R. Kulur, Marcos A.S. Fonseca, Se-Jin Yoon, Chunyang Dong, Rebecca J. Levy, Min-Yin Li, Enrique Sosa, Abel Bermudez, Delaney Wood, Isabelle Liu, Amander T. Clark, Anne Brunet, Sharon Pitteri, Kevin Kelley, Maria Barna, Sergiu P. Paşca

## Abstract

Human pluripotent stem cell-derived models are increasingly used to recapitulate aspects of human corticogenesis, including the emergence of diverse cortical cell types and their progression through developmental states. These systems offer an opportunity to dissect the molecular mechanisms regulating transitions between cell states, but methods for perturbing them systematically at scale remain limited. Here, we developed a semi-automated culture and high-throughput arrayed screening platform for human cortical organoids. Using a library of 1,280 small molecules designed to systematically target major signaling and regulatory pathways, we screened for modulators of *EOMES*, which encodes for the transcription factor TBR2, and identified Emetine, an inhibitor of translation elongation, as the top hit. The effect of Emetine on cortical cell-state progression was reproduced across human pluripotent stem cell lines, non-human primate cortical organoids, and primary human cortical tissue but not broadly across translation inhibitors. Moreover, transient exposure resulted in a shift from dividing radial glia towards cortical glutamatergic neurons. Mechanistically, nascent-proteome profiling revealed an early and coordinated remodeling of newly translated proteins that preceded detectable transcriptional changes, placing translational regulation upstream of the ensuing cell-state transition. Together, these findings highlight a role for specific translational inhibitors in promoting cortical cell-state progression and establish a scalable platform for systematic perturbation and discovery in human neural organoids.

## Introduction

Recent advances in human stem cell-derived models are making it possible to reconstruct aspects of the developmental sequence of corticogenesis, from progenitor expansion to the emergence of diverse cortical cell types^1-4^. Regionalized neural organoids, including human cortical organoids (hCO) that resemble dorsal forebrain, generate many of the progenitor and neuronal populations found in the developing human cerebral cortex and recapitulate aspects of their temporal emergence, progressing from radial glia (RG) through intermediate progenitor cell (IPC) states to the sequential generation of deep- and upper-layer cortical neurons^5,6^, followed by the emergence of glial cells^1,7^. Defining the mechanisms that control progression through these states is important for understanding how developmental trajectories are altered in disease^8^ and how these programs have changed during primate and human evolution^9^. Human stem cell-derived models provide an opportunity to interrogate these developmental transitions experimentally. Although pooled genetic screens have begun to demonstrate the potential of systematic perturbation in neural organoids^10-12^, their use for arrayed screening remains limited. Neural organoid cultures require prolonged differentiation and repeated media exchanges, and a combination of cost, variability, and the limited availability of scalable quantitative readouts has made it difficult to perform arrayed screens^13-15^. As a result, the ability of these models to resolve developmental cell states has advanced more rapidly than our ability to systematically perturb them and identify the molecular mechanisms controlling cell state transitions. Understanding these transitions may be particularly important in the human cerebral cortex, where progenitor populations and their developmental trajectories have undergone substantial evolutionary changes, placing additional demands on the mechanisms that coordinate proliferation and differentiation^16,17^. Among these, *EOMES* (also known as TBR2) is expressed across transient states, including early Cajal–Retzius (CR) cells^18,19^ and IPCs (also known as transient amplifying progenitors) that contribute to the generation of cortical glutamatergic neurons^20^. The expansion of cortical progenitor populations has been considered an important feature of primate corticogenesis^21^, while perturbations in their proliferation or differentiation are thought to contribute to neurodevelopmental disease^22^. Here, we developed a semi-automated, low-cost approach that enables arrayed perturbation of hCO at scale. We used this platform to systematically survey major signaling and regulatory pathways for their ability to alter progression in early cortical states. Using *EOMES* as a quantifiable readout for IPCs, this screen identified Emetine, a translation elongation inhibitor, as a potent regulator of the transition from proliferative progenitors toward differentiated neurons. Follow-up experiments in primate cortical organoids and human primary tissue slices showed that this effect reflects a change in developmental progression. Time-resolved transcriptomic and nascent proteome profiling revealed that changes in newly synthesized proteins emerge before substantial transcriptional changes, suggesting that modulating protein synthesis can shape subsequent changes in cell identity. Together, these studies establish a framework for perturbing developmental trajectories directly in human organoid models and that modulating translational control represents a means to regulate the pace of early corticogenesis.

## Results

### Generation of a scalable cortical organoid culturing and arrayed screening platform

To establish a user-friendly and low-cost culture and screening platform, we adapted our previous human cortical organoid protocol^15^ to a 96-well format with incorporation of a washer-dispenser system for semi-automated media exchange (Fig. 1a and Fig. S1a). Organoids generated in this format grew reproducibly over time across 50 days in culture (Fig. 1b, Fig. 1c and Fig. S1b). We next benchmarked the 96-well protocol against our bulk hCO protocol^15^ (Fig. S1c) and found no differences in culture quality as seen by quality control markers observed by qPCR, supporting the suitability of the high-throughput format for systematic perturbation studies.

**Fig. 1:**
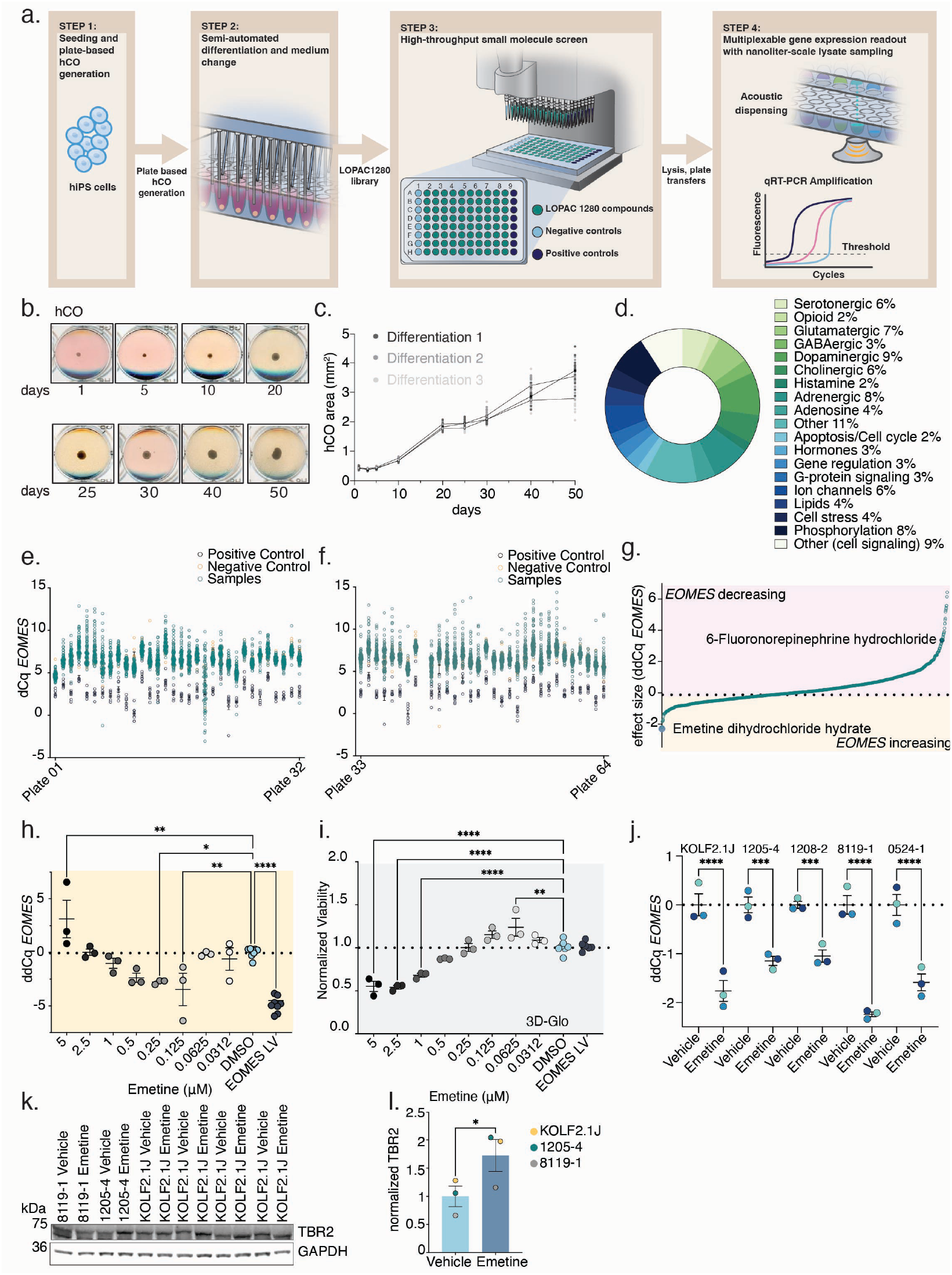
Establishment of a high-throughput neural organoid culture platform and screening of LOPAC1280 library for regulators of *EOMES* expression. a) Schematic depicting the generation of 96-well based, semi-automated organoid cultures and workflow of the high-throughput screening platform. b) Representative images of hCO growth in a 96-well plate over 50 days of neural differentiation. **c)** Quantification of organoid growth in 96-well plates over 50 days of neural differentiation (n = 46 to 48 organoids from 3 differentiations). Each dot represents an individual organoid; data are shown as mean ± SEM. **d)** Constitution of the target classes of the LOPAC1280 small molecule library. **e** and **f)** *EOMES* expression across the LOPAC1280 small molecule screen, shown as dCq values for individual wells measured by μqPCR. Samples and controls are grouped by screening plate along the x axis. Overexpression of *EOMES* by a lentivirus is used as a positive control. e) Low concentration screening subset (50 nM; plates 1–32). f) High-concentration subset (1 μM; plates 33–64). **g)** Effect size of LOPAC1280 compounds on *EOMES* expression at 1 μM. ddCq values are shown for each well relative to negative controls and were used for hit selection. Emetine dihydrochloride hydrate, which increased *EOMES* expression, and 6-Fluoronorepinephrine hydrochloride that decreased *EOMES* expression are highlighted by enlarged symbols. **h)** Dose response validation of increase of *EOMES* expression by Emetine in the secondary screen. Each dot represents an individual well quantified by μqPCR and reported as ddCq relative to DMSO controls. Overexpression of *EOMES* is used as a positive control for the assay. Ordinary one-way ANOVA followed by Dunnett’s multiple comparisons test; F(9,30) = 14.62, P = <0.0001; for 5 μM **P = 0.0079, for 0.25 μM *P = 0.0127, for 0.125 μM **P = 0.0026, for EOMES LV ****P = <0.0001. Data are represented as mean ± SEM. **i)** Dose response assessment of hCO viability following Emetine treatment in the secondary screen. Each dot represents an individual well quantified by 3D-Glo viability assay normalized to DMSO-treated controls. Ordinary one-way ANOVA followed by Dunnett’s multiple comparisons test; F(9,27) = 31.03, P = <0.0001; for 5 μM, 2.5 μM and 1 μM ****P= <0.0001, for 0.0625 μM **P = 0.0020. Data are represented as mean ± SEM. **j)** Validation of Emetine related increase of *EOMES* expression across hCO derived from five different hiPS cell lines. Each dot represents an independent differentiation (n=3 differentiations per line). EOMES expression was quantified by qRT-PCR and effect size is reported as ddCq relative to Vehicle-treated controls. Ordinary one-way ANOVA followed by Dunnett’s multiple comparisons test; F(9,20) = 29.18, P = <0.0001; KOLF2.1J ****P = <0.0001, 1205-4 ***P = 0.0004, 1208-2 ***P = 0.0010, 8119-1 ****P = <0.0001, 0524-1 ****P = <0.0001. Data are represented as mean ± SEM. **k)** Representative immunoblot of TBR2 in hCO following Emetine treatment. Each treatment pair represents an independent differentiation (n= 6) from 3 hiPS cell line derived hCO. GAPDH was used as a loading control. **l)** Quantification of western blots for TBR2 expression following Emetine treatment of hCO. Paired t-test was performed with all 3 lines after averaging of KOLF2.1J to normalize for differing differentiation numbers across the samples; P = 0.0368. Data are represented as mean ± SEM.

To establish a screening readout, we focused on *EOMES*, which marks early cortical progenitor and neuronal populations arising between RG expansion and early neuronal differentiation^23^. This enabled us to capture perturbations affecting progenitor progression as well as downstream consequences for early neuronal differentiation. To quantify *EOMES* expression in a scalable manner, we implemented one-step micro-volume qPCR (µqPCR) using nanoliter-scale lysate sampling and acoustic dispensing (Fig. 1a and Fig. S1d). We compared the one-step qPCR approach with conventional two-step qPCR across multiple human pluripotent stem cell lines (hiPS) from day 25 to day 50 of neural differentiation (Fig. S1e and Fig. S1f). In KOLF2.1J-derived hCO, *EOMES* expression peaked around day 30 of differentiation; therefore, we proceeded with this line for the primary screen. To enable biological interpretation of candidate regulators emerging from this platform, we selected the target-annotated LOPAC1280 small-molecule library, targeting all major cell signaling pathways and neurotransmission (Fig. 1d) to be applied across three media changes from day 25 to 31 of differentiation and *EOMES* µqPCR (Fig. 1a and Fig. S1d) as the primary screening conditions. Together, these optimizations allowed scaling of hCO differentiation into an arrayed perturbation platform enabling quantitative molecular screening across thousands of conditions.

### A high-throughput small molecule hCO screen for regulators of *EOMES* expression

Using this platform, we screened a total of 6,144 hCO corresponding to duplicates across two concentrations, 50 nM and 1 µM, following 6 days of compound exposure from 25 to 31 days of neural differentiation (Fig. S1d). Duplicates were reformatted to be on separate plates and each screening plate contained lentiviral *EOMES* overexpression as a positive control and DMSO as a negative control (n=8 wells on each plate for positive and negative control respectively, Fig. 1a, Step 3) for the µqPCR readout. Across screened plates, positive and negative controls were well separated, supporting assay robustness (Fig. 1e and Fig. 1f).

We ranked hits by effect size relative to DMSO-treated controls across replicate wells (Fig. 1g). Among all conditions tested, emetine dihydrochloride hydrate, hereafter referred to as Emetine, emerged as the strongest candidate for increasing *EOMES* expression (Fig. 1g). The screen also identified the serotonin receptor agonist (±)-8-hydroxy-DPAT hydrobromide as a candidate for reducing *EOMES* expression, consistent with a broader role for serotonergic signaling in human cortical progenitor regulation^24^. To address whether additional compounds targeting neuromodulatory signaling could alter *EOMES* expression, we selected the adrenergic agonist 6-fluoronorepinephrine hydrochloride and the cholecystokinin receptor antagonist Proglumide from the top 10 candidates for follow-up (Table S1). We performed a secondary dose-response screen using an eight-point serial dilution series, with parallel assessment of cell viability using 3D-Glo, using ATP content as a proxy (Fig. 1h, i and Fig. S1g–j). In the secondary screen, Norepinephrine and Proglumide showed no clear effect in the conditions tested (Fig. S1g and i). The top candidate from the primary screen, Emetine, showed a robust increase of *EOMES* expression at 250 nM (Fig. 1h), without compromising viability. Consistent with its known activity as a translation elongation inhibitor that directly binds to the E-site of the 40S in the assembled 80S ribosome^25^, Emetine reduced viability at higher concentrations (Fig. 1i). We therefore selected 250 nM Emetine as the working concentration for subsequent experiments. Together, these results demonstrate the utility of the unbiased hCO screening platform to identify compounds that alter *EOMES*, highlighting Emetine for its dose-dependent increase in *EOMES* levels within a viability-compatible concentration range.

We next tested whether an increase in *EOMES* expression following Emetine treatment was reproducible across independent hiPS cell lines and differentiations. Across all five tested hiPS cell line derived hCO, 250 nM Emetine treatment from day 25 to day 31 of neural differentiation (hereafter referred to as Emetine treatment) consistently increased *EOMES* expression, indicating that the effect was not restricted to the screening line or to a single differentiation batch (Fig. 1j). To rule out the possibility that the change in *EOMES* mRNA levels diverged from a change in protein levels, we performed immunoblotting following Emetine treatment and observed an increase in TBR2 signal (Fig. 1k and Fig. 1l), demonstrating that the transcriptional increase detected by µqPCR was reflected at the protein level.

### Emetine treatment across a range of human and non-human primate model systems

Since the emergence and diversification of cortical progenitor populations, including *EOMES*^+^ IPCs have been linked to an evolutionary increase in neocortical size and complexity^17,20,26^, we questioned whether we could recapitulate this effect in a non-human primate organoid system. We generated macaque cortical organoids (mqCO) (Fig. 2a) from two independent macaque iPS (mqiPS) cell lines^27,28^ across four differentiations and treated them with Emetine. Emetine increased *EOMES* expression across macaque lines and differentiation batches at 31 days of neural differentiation (Fig. 2b), indicating that the response extends beyond hCO and is present in a non-human primate cortical model.

**Fig. 2:**
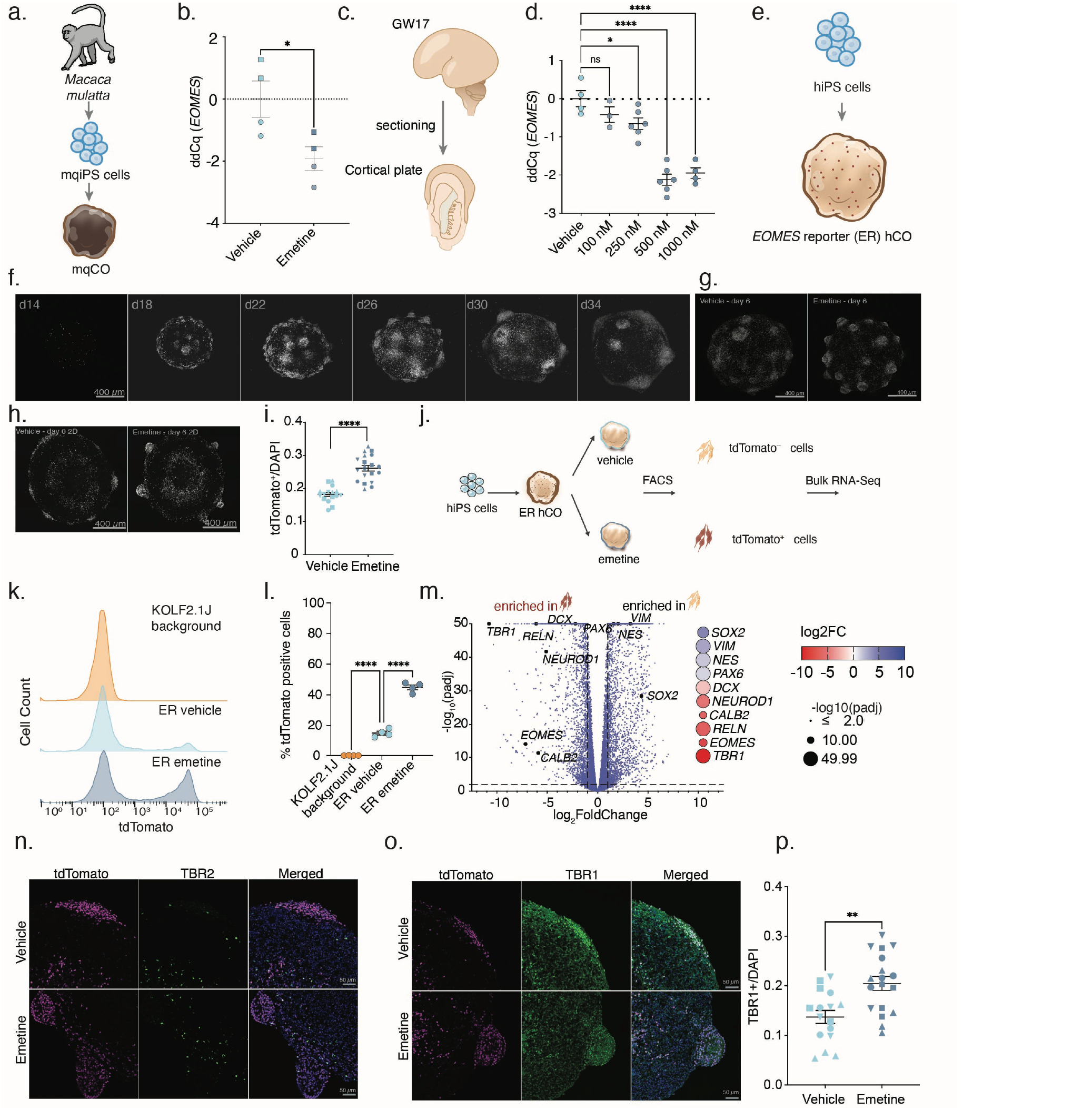
Validation of increase in *EOMES* expression following Emetine treatment across human and non-human primate cortical models. **a)** Schematic of macaque cortical organoid (mqCO) generation from *Macaca mulatta* riPSC05 and riPSC89 iPS cell lines. **b)** qRT-PCR of EOMES expression levels of mqCO following treatment with 250 nM Emetine from day 25-31 of neural induction. n=4 independent differentiations and two mqiPS cell lines; square symbol indicates riPSC05, circle symbol indicates riPSC89. EOMES expression was quantified by qRT-PCR and effect size is reported as ddCq relative to Vehicle-treated controls. Data are represented as mean ± SEM. Data were statistically assessed with two-tailed t-test; *P = 0.0326. **c)** Schematic of *ex vivo* sliced human primary cortical cultures at GW17. **d)** Emetine treatment of human primary cortical tissue. Expression of EOMES was quantified by qRT-PCR and effect size is reported as ddCq to Vehicle-treated controls. Each dot reports an independently cultured slice preparation. Ordinary one-way ANOVA followed by Dunnett’s multiple comparisons test; F(4,18) = 32.38, P = <0.0001; for 250 nM *P = 0.0355, for 500 nM ****P = <0.0001, for 1000 nM ****P = <0.0001. Data are represented as mean ± SEM. **e)** Schematic of the differentiation of EOMES reporter (ER) to hCO. **f)** Representative z-projected live confocal images of tdTomato signal in ER hCO in a 96-well plate from d14 to d34 of neural differentiation. Scale bar = 400 μm. **g)** Representative z-projected live confocal images of Vehicle and Emetine treated ER hCO. Scale bar = 400 μm. **h)** Representative fluorescence images of embedded and cryosectioned ER hCO following Vehicle and Emetine treatment for 6 days for tdTomato signal. **i)** Quantification of tdTomato^+^/DAPI signal following Emetine treatment (n = 36 organoids from 4 differentiations); Welch’s two-tailed t-test, ****P= <0.0001. Data are represented as mean ± SEM. **j)** Schematic detailing derivation of ER hCO and Emetine treatment, followed by FACS isolation of tdTomato^+^ and tdTomato^-^ cells for bulk RNA-Seq at 31 days of neural differentiation. **k)** Representative FACS plot of ER hCO following vehicle and emetine treatment. Parental KOLF2.1J-derived hCO were used as a fluorescence background control. **l)** Quantification of sorted tdTomato^+^ cells from KOLF2.1J hCO, ER Emetine and Vehicle treated hCO (n = 4 independent differentiations). Ordinary one-way ANOVA followed by Dunnett’s multiple comparisons test; F(2,9) = 376.8, P = <0.0001; for ER Vehicle vs. KOLF2.1J background ****P = <0.0001, for ER Emetine vs ER Vehicle ****P = <0.0001. Data are represented as mean ± SEM. **m)** Differential gene expression between FACS isolated tdTomato+ and tdTomato− cells from ER hCO treated with Vehicle. The volcano plot shows log2 fold change and −log10(adjusted P value) for each gene. Negative log2 fold-change values indicate genes enriched in tdTomato^+^ cells, whereas positive values indicate genes enriched in tdTomato− cells. Selected neural progenitor and neuronal lineage markers are highlighted. Color indicates log2 fold change, and symbol size indicates −log10(adjusted P value) (n= 4 independent differentiations for each condition). **n)** Representative immunocytochemistry (TBR2) and fluorescence (tdTomato) images of Vehicle and Emetine treated ER hCO at 31 days of differentiation. tdTomato is shown in magenta, TBR2 in green and DAPI in blue. Scale bar = 50 μm. **o)** Representative immunocytochemistry (TBR1) and fluorescence (tdTomato) images of Vehicle and Emetine treated ER hCO at 31 days of differentiation. tdTomato is shown in magenta, TBR1 in green and DAPI in blue. Scale bar = 50 μm. **p)** Quantification of TBR1^+^/DAPI signal following Emetine treatment (n = 34 organoids from 4 differentiations and 2 hiPS cell lines, shapes indicate differentiations); Welch’s two-tailed t-test, **P= 0.0013. Data are represented as mean ± SEM.

We next asked whether the effect could be observed in human primary tissue. We generated cortical slice cultures from gestational week (GW) 17 (Fig. 2c) and treated them with increasing concentrations of Emetine, ranging from 100 to 1000 nM for 6 days. Emetine increased *EOMES* signal in a dose-dependent manner in human primary cortical tissue (Fig. 2d).

To enable visualization and cellular quantification of *EOMES*-associated cortical states, we generated an *EOMES* reporter line in the KOLF2.1J hiPS cell background by introducing an NLS-tdTomato cassette into the endogenous *EOMES* locus (hereafter ER; Fig. S2a). We next differentiated the ER line into hCO (Fig. 2e) and monitored reporter fluorescence longitudinally and observed detectable tdTomato signal from approximately day 14 of neural differentiation which continuously increased until reaching a maximum around day 25 and slowly decreasing thereafter (Fig. 2f), paralleling the developmental profile of endogenous *EOMES* expression measured by qRT-PCR. Immunostaining for endogenous TBR2 at day 20 of neural differentiation showed a 75% average overlap between tdTomato and TBR2 signal (Fig. S2b and Fig. S2c), supporting concordance between reporter activity and endogenous *EOMES*/TBR2 expression. We then confirmed that ER hCO retained the response to Emetine observed in parental KOLF2.1J hiPS cell line-derived hCO. Across five independent differentiations, Emetine treatment reproducibly increased *EOMES* expression measured by qRT-PCR (Fig. S2d).

In accordance with the qRT-PCR and immunoblotting results, Emetine treatment increased tdTomato reporter signal in ER hCO. Whole-organoid live imaging revealed an expanded reporter-positive domain following Emetine exposure (Fig. 2g, Movie S1), and quantitative analysis of cryosectioned hCO confirmed an increase in tdTomato+ signal (Fig. 2h, i).

We next used the ER line to quantify this response at single cell resolution and to define the molecular states captured by the reporter. Vehicle and Emetine-treated ER hCO were dissociated at day 31 and analyzed by fluorescence-activated cell sorting (FACS) based on tdTomato signal (Fig. 2j). Emetine increased the fraction of tdTomato^+^ cells across four independent differentiations and cell sorting experiments (Fig. 2k, l and Fig. S2e), demonstrating that the increase detected by bulk molecular assays was accompanied by an expansion on the single cell level. In parallel, tdTomato^+^ and tdTomato-populations were isolated for bulk RNA-Seq (Fig. 2j). Transcriptomic profiling confirmed enrichment of *EOMES* in the tdTomato^+^ fraction (Fig. 2m) by differential gene expression (DEG). In addition to *EOMES*, tdTomato^+^ cells were enriched for RELN and CALB2, associated with early CR-like states, together with early neuronal genes including TBR1, DCX, and NEUROD1 (Fig. 2m). In contrast, tdTomato-cells were enriched for RG markers including PAX6, NES, VIM, and SOX2. These results indicate that ER fluorescence at this developmental stage captures a heterogeneous early cortical population rather than a single transcriptionally defined progenitor population. Intrigued by these results, we performed TBR2 (Fig. 2n, Fig. S2f) immunostaining on the Vehicle and Emetine treated ER hCO on day 31 of differentiation and observed that colocalization of the tdTomato and TBR2 signal had reduced dramatically to an average of 3.7% (Fig. S2g) and that expanded reporter-positive domains under Emetine treatment were not positive for the TBR2 staining. In contrast, these domains were positive for TBR1 staining (Fig. 2o and Fig. S2h, lower panels) and we found an increase in TBR1 positive cells in Emetine treated hCO (Fig. 2p). In addition, the increase in TBR1 was confirmed in human primary slice cultures treated with Emetine (Fig. S2i). These results suggest first that Emetine may result in a fate-shift of progenitor populations towards newborn neurons and that the ER hCO line retains the tdTomato signal beyond the transient endogenous *EOMES* expression window, providing a short-to medium-term record of cells that recently occupied an *EOMES*-expressing state, as previously reported^29^. Moreover, the heterogeneity observed in the ER hCO is consistent with the transient and developmentally dynamic expression of *EOMES*. Even though *EOMES*/TBR2 is prominently expressed by IPCs, *EOMES* expression has also been described in early preplate and CR populations and in cells transitioning toward neuronal differentiation, in primary first-trimester human brain and longitudinal cortical organoid atlases^2,30^, and human stem cell-derived cortical cultures contain RELN-positive presumptive CR cells that frequently express *EOMES*^19,31^.

Following characterization of the ER line we sought to understand how Emetine altered the molecular state of the tdTomato^+^ cells. Principal component analysis of bulk RNA-Seq profiles showed clear separation of Vehicle and Emetine-treated tdTomato^+^ samples across four independent differentiations (Fig. S2j), indicating a reproducible transcriptional response to Emetine. DEG between Emetine and Vehicle treatment identified a concordant set of strongly altered genes across differentiation batches, illustrated by the top differentially expressed genes (Fig. S2k). Gene Ontology (GO) analysis of increased DEG following Emetine treatment revealed enrichment of biological processes associated with translational regulation and protein synthesis (Fig. S2l), which may reflect transcriptional feedback onto translation consistent with the known activity of Emetine on translation elongation^32^. Together, these data show that Emetine reproducibly increases EOMES-associated readouts across hCO, primary human fetal cortex, and mqCO.

### Single cell profiling of Emetine treated hCO

Seeing an increase in TBR1^+^ cells, we wanted to investigate the changes to the cortical cell type composition in an unbiased manner and therefore performed scRNA-Seq of day 31 hCO derived from three hiPS cell lines across two differentiations following Vehicle or Emetine treatment (Fig. S3a). We found that RG, dividing progenitors, IPC, CR, and early born neurons were consistently represented across cell lines and differentiations (Fig. S3b). Consistent with the reporter-based FACS and bulk RNA-Seq results, Emetine increased the proportion of *EOMES*-expressing cells across lines and differentiations (Fig. 3a, b and Fig. S3c). Unsupervised clustering and marker-gene analysis identified the expected major early cortical populations, including RG, IPC, CR, and early newborn neurons (Fig. 3c, d and Fig. S3e).

**Fig. 3:**
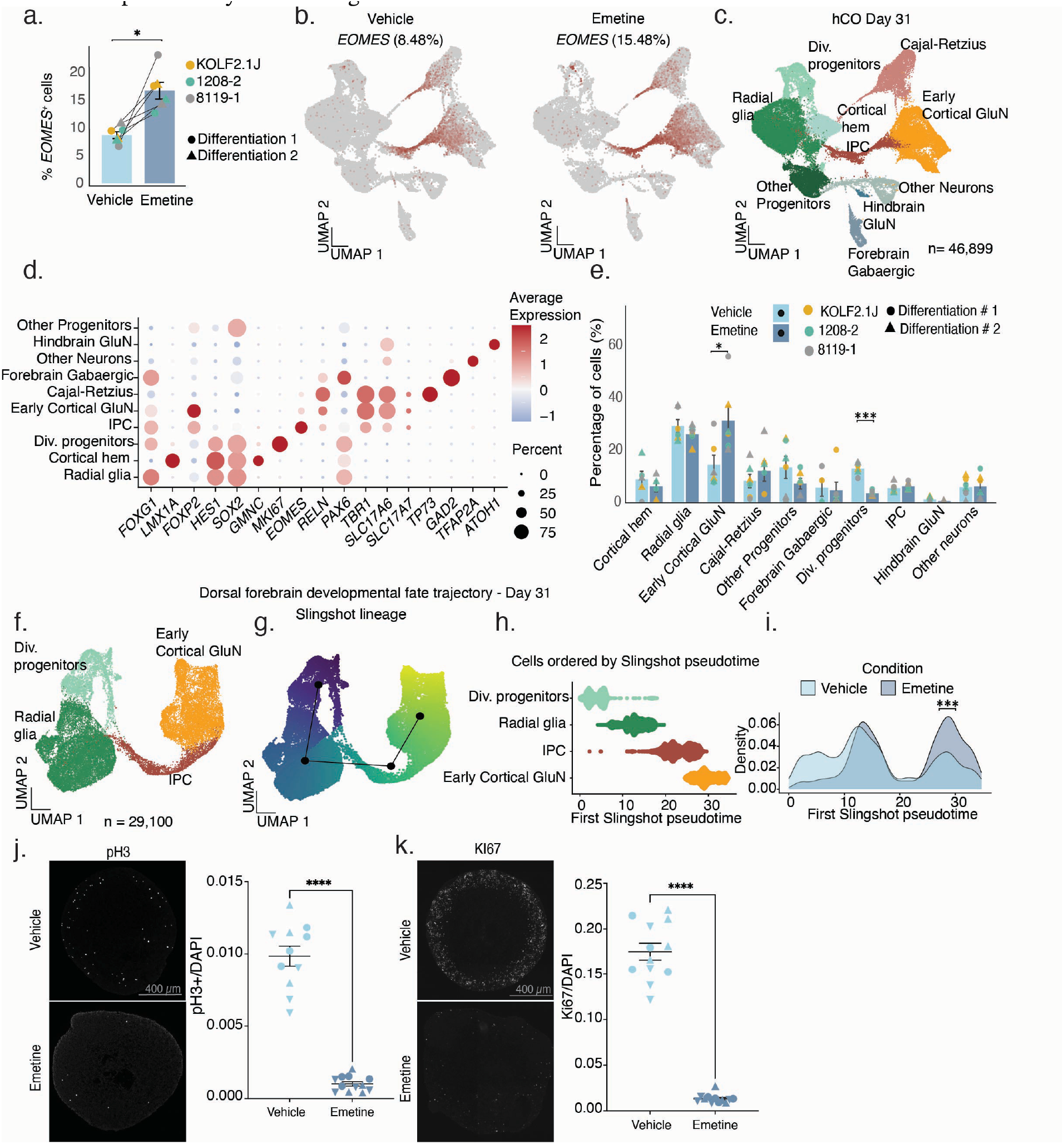
Single cell and molecular profiling of Emetine treated hCO at day 31 of neural differentiation. **a)** Percentage of *EOMES*^*+*^ cells from Vehicle or Emetine treated hCO (n= 3 hiPS cell lines and 2 differentiations for each condition). Paired t-test; *P = 0.0104. **b)** Uniform manifold approximation and projection (UMAP) visualization of *EOMES*^*+*^ cells in Vehicle (8.48%) or Emetine treatment (15.48%). Positive cells shown in crimson all other cells are shown in grey. **c)** UMAP visualization of single-cell gene expression of hCO at day 31 following Vehicle or Emetine treatment and cell type annotation (n = 46,899 cells from three hiPS cell lines and two differentiations for each condition). **d)** Dot plots of marker gene expression in Vehicle and Emetine treated hCO. The size of the circles represents the percentage of cells expressing the marker and color of the dots represent scaled gene expression values. **e)** Bar plot of cells that are annotated to each cluster, expressed as percentage of cells from total cells for Vehicle and Emetine treatment conditions. Linear mixed model considering cell line and differentiation as random effects, Benjamini-Hochberg adjusted ***P = < 9.5 × 10^−07^, *P= 0.0461. **f)** UMAP for subset of cells of dorsal cortical lineage progression (n= 29,100 cells from 3 hiPS cell lines and 2 differentiations). **g)** Visualization of Slingshot lineage progression of dorsal forebrain cells with associated pseudotime distributions along the lineage trajectory. **h)** Dorsal forebrain cell types ordered by pseudotime. **i)** Per-condition kernel density estimation of the pseudotime distributions. Permutation test P-value = 0.0005. **j)** Representative immunocytochemistry images of pH3 staining in embedded and cryosectioned KOLF2.1J-derived hCO following Vehicle and Emetine treatment. Quantification of pH3^+^ cells/DAPI signal (n = 23 organoids from 3 differentiations, shapes indicate differentiations); Welch’s two-tailed t-test, ****P = <0.0001. Data are represented as mean ± SEM. **k)** Representative immunocyto-chemistry images of KI67 staining in embedded and cryosectioned KOLF2.1J-derived hCO following Vehicle and Emetine treatment. Quantification of % of KI67^+^ cells/DAPI signal (n = 24 organoids from 3 differentiations, shapes indicate differentiations), Welch’s two-tailed t-test, ****P = <0.0001. Data are represented as mean ± SEM.

We next asked whether Emetine altered cortical cell identity or induced aberrant transcriptional states. Emetine-treated cells remained distributed within the same transcriptional cell-state landscape as Vehicle-treated cells, without the emergence of treatment-specific clusters or broad loss of cortical identity (Fig. S3d). Thus, Emetine altered the representation of existing developmental states. Quantification of cell-state proportions revealed a reduction in dividing progenitor populations following Emetine treatment, accompanied by increased representation of early newborn neuronal states (Fig. 3e). We therefore examined the dorsal cortical developmental trajectory using Slingshot pseudotime analysis (Fig. 3f-h). Emetine-treated cells were significantly shifted toward later positions along the inferred radial glia-to-newborn-neuron trajectory (Fig. 3i), consistent with enhanced progression from proliferative progenitor toward early neuronal states. Immunocytochemistry for proliferative markers pH3 and KI67 independently confirmed the reduction in dividing progenitor populations (Fig. 3j, k). Together with the increase in TBR1^+^ cells, these findings indicate that Emetine shifts the balance of early cortical states away from proliferative radial glia and toward early-born neuronal populations.

We next asked whether transient Emetine exposure produced persistent alterations in subsequent stages of cortical development. hCO derived from four independent hiPS cell lines were exposed to Emetine from day 25 to day 31, returned to standard culture conditions, then profiled by scRNA-Seq at day 50 (Fig. S4a). The data showed broad representation of RG, dividing progenitors, IPCs, CR cells, and excitatory neurons across hiPS cell lines (Fig. S4b). Clustering and marker-gene analysis identified the expected cortical cell states (Fig. S4c-e), and prior Emetine exposure did not result in loss of cortical identity or emergence of non-cortical or treatment-specific populations (Fig. S4f). Analysis of dorsal cortical developmental trajectories further revealed no significant difference in pseudotime progression between previously treated and control organoids at day 50 (Fig. S4g-j). Thus, the shift observed immediately following Emetine exposure was not associated with a detectable persistent disruption of subsequent cortical developmental progression. Together, these results show that Emetine exposure is associated with a transient shift in the early cortical developmental landscape from dividing progenitor states towards early neuronal states.

### Characterization of translation inhibition and stress response pathways following Emetine exposure

To ask whether Emetine’s effect on neural progenitor progression was a general consequence of suppressing protein synthesis, we tested eight compounds acting at different points in the translation machinery (Fig. 4a). These included 4EGI-1, which disrupts the eIF4E-eIF4G interaction required for cap-dependent translation initiation; INK128, which inhibits upstream mTORC1/2 signaling^33^; and Homoharringtonine, which blocks the initial elongation step. Along with these we also tested Puromycin, which causes premature release of nascent polypeptide chains; Cycloheximide and Cryptopleurine, which inhibit ribosomal translocation; Anisomycin, which inhibits peptide-bond formation; and Didemnin B, which inhibits eEF1A-dependent aminoacyl-tRNA accommodation^34^. For each compound, we performed an eight-point dose-response analysis following treatment from 25-31 days of neural differentiation with parallel measurements of cell viability and *EOMES* expression. All compounds showed reduced viability at higher concentrations, allowing us to define concentration ranges in which *EOMES* regulation could be assessed in the absence of overt toxicity (Fig. 4b, d, f, h and Fig. S5a, c, e, g). Within these ranges, Puromycin (Fig. 4c), Cycloheximide (Fig. 4e), 4EGI-1 (Fig. S5b), Anisomycin (Fig. S5d), and Homoharringtonine (Fig. S5h) did not increase *EOMES* expression. INK128 produced a modest decrease in *EOMES* at 25 nM (Fig. S5f). By contrast, only two of the eight perturbations increased *EOMES*: Cryptopleurine (Fig. 4g), which binds a ribosomal site overlapping with Emetine^32,35^, and Didemnin B (Fig. 4i). Thus, increased EOMES was not a general consequence of suppressing protein synthesis but may depend on other features of elongation perturbation, including selectivity of the mode of perturbation^36^, or the persistence of translational inhibition.

**Fig. 4:**
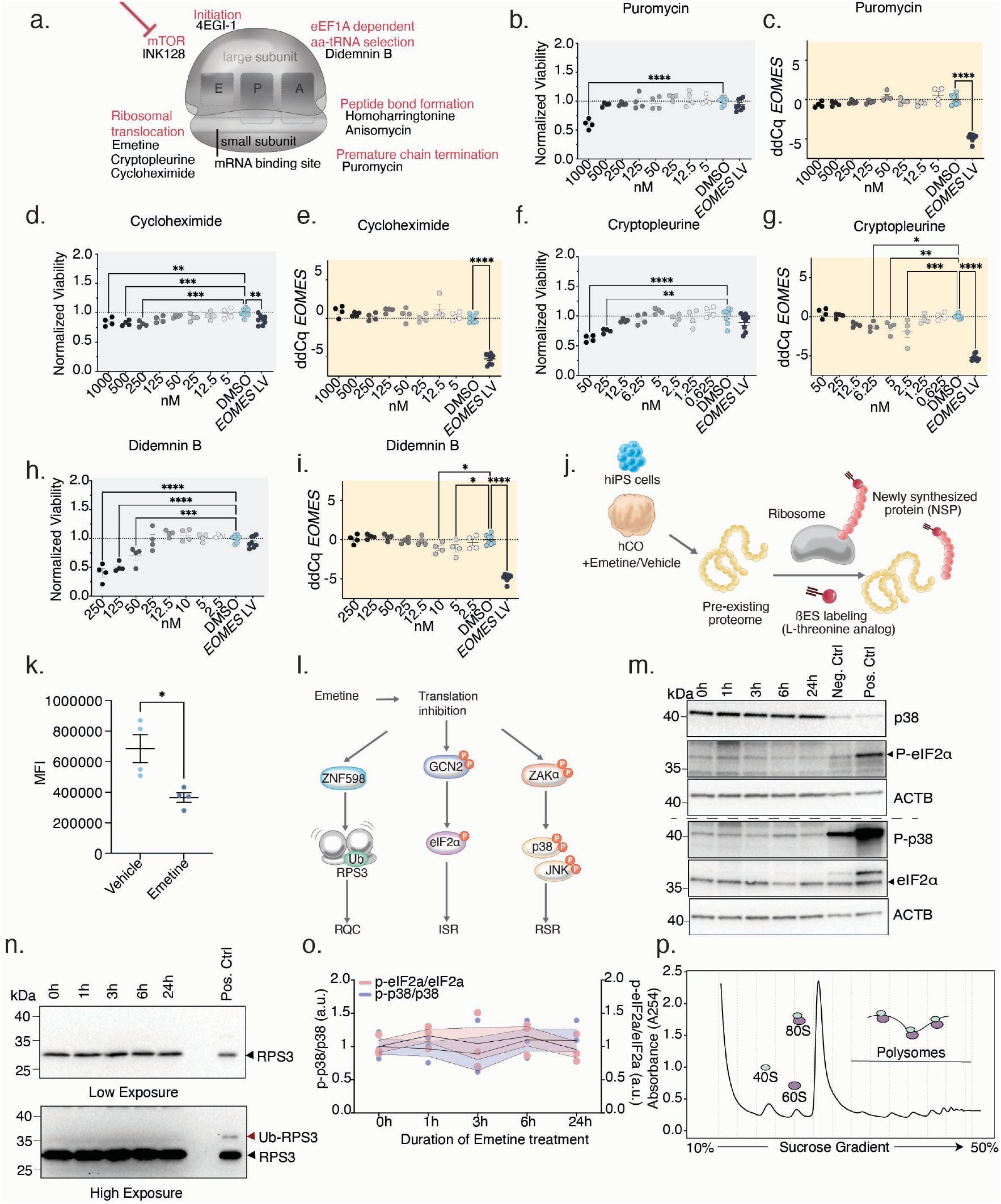
Characterization of translation inhibition, stress response pathways following Emetine exposure and basal polysome profile of hCO. **a)** Schematic detailing diversity of the translation inhibitors tested. **b)** Dose response assessment of hCO viability following Puromycin treatment in the translation inhibitor screen. Each dot represents an individual well quantified by 3D-Glo viability assay normalized to DMSO-treated controls. Ordinary one-way ANOVA followed by Dunnett’s multiple comparisons test; F(9,38) = 9.319, P = <0.0001; for 1000 nM ****P= <0.0001. Data are represented as mean ± SEM. **c)** Dose response of *EOMES* expression by Puromycin in the translation inhibitor screen. Each dot represents an individual well quantified by μqPCR and reported as ddCq relative to DMSO controls. *EOMES* overexpression is used as a positive control for the assay. Ordinary one-way ANOVA followed by Dunnett’s multiple comparisons test; F(9,38) = 65.02, P = <0.0001; *EOMES* LV ****P = <0.0001. Data are represented as mean ± SEM. **d)** Dose response assessment of hCO viability following Cycloheximide treatment in the translation inhibitor screen. Each dot represents an individual well quantified by 3D-Glo viability assay normalized to DMSO-treated controls. Ordinary one-way ANOVA followed by Dunnett’s multiple comparisons test; F(9,38) = 5.210, P = 0.0001; for 1000 nM **P= 0.0039, for 500 nM ***P=0.0007, for 250 nM ***P=0.0002, for *EOMES* LV **P=0.0072. Data are represented as mean ± SEM. **e)** Dose response of *EOMES* expression by Cycloheximide in the translation inhibitor screen. Each dot represents an individual well quantified by μqPCR and reported as ddCq relative to DMSO controls. *EOMES* overexpression is used as a positive control for the assay. Ordinary one-way ANOVA followed by Dunnett’s multiple comparisons test; F(9,38) = 54.01, P = <0.0001; *EOMES* LV ****P = <0.0001. Data are represented as mean ± SEM. **f)** Dose response assessment of hCO viability following Cryptopleurine treatment in the translation inhibitor screen. Each dot represents an individual well quantified by 3D-Glo viability assay normalized to DMSO-treated controls. Ordinary one-way ANOVA followed by Dunnett’s multiple comparisons test; F(9,38) = 7.619, P = <0.0001; for 50 nM ****P= <0.0001, for 25 nM **P= 0.0027. Data are represented as mean ± SEM. **g)** Dose response of *EOMES* expression by Cryptopleurine in the translation inhibitor screen. Each dot represents an individual well quantified by μqPCR and reported as ddCq relative to DMSO controls. *EOMES* overexpression is used as a positive control for the assay. Ordinary one-way ANOVA followed by Dunnett’s multiple comparisons test; F(9,38) = 47.73, P = <0.0001; for 6.25 nM *P= 0.0156, for 5 nM **P= 0.0064, for 2.5 nM ***P= 0.0001, for *EOMES* LV ****P = <0.0001. Data are represented as mean ± SEM. **h)** Dose response assessment of hCO viability following Didemnin B treatment in the translation inhibitor screen. Each dot represents an individual well quantified by 3D-Glo viability assay normalized to DMSO-treated controls. Ordinary one-way ANOVA followed by Dunnett’s multiple comparisons test; F(9,38) = 23.27, P = <0.0001; for 250 nM ****P= <0.0001, for 125 nM ****P= <0.0001, for 50 nM ***P=0.0003. Data are represented as mean ± SEM. **i)** Dose response of *EOMES* expression by Didemnin B in the translation inhibitor screen. Each dot represents an individual well quantified by μqPCR and reported as ddCq relative to DMSO controls. *EOMES* overexpression is used as a positive control for the assay. Ordinary one-way ANOVA followed by Dunnett’s multiple comparisons test; F(9,38) = 46.98, P = <0.0001. for 10 nM *P= 0.0435, for 5 nM *P= 0.0371, for *EOMES* LV ****P = <0.0001. Data are represented as mean ± SEM. **j)** Schematic of β-Ethynylserine (βES) workflow for nascently synthesized peptide labeling. **k)** Quantification of βES labeled ER derived hCO dissociated cells following Emetine or Vehicle treatment by FACS analysis (n= 4 differentiations). Welch’s two-tailed t-test, *P= 0.0334. Data are represented as mean ± SEM. **l)** Schematic depicting canonical stress-response and quality-control pathways engaged by translational perturbation. ISR, integrated stress response; RSR, ribotoxic stress response; RQC, ribosome-associated quality control. **m)** Representative immunoblot of phosphorylated EIF2α (pEIF2α) and total EIF2α in hCO following Emetine treatment time course from 0h to 24h for assessing ISR activation and representative immunoblot of phosphorylated p38 (p-p38) and total p38 in hCO following Emetine treatment time course from 0h to 24h for assessing RSR activation. **n)** Representative immunoblot of ubiquitylated RPS3 in hCO following Emetine treatment time course from 1 hour to 24 hours for assessing RQC activation on a sucrose cushion. **o)** Quantification of western blots for ISR, RSR activation from panels m following Emetine treatment of hCO. For each immunoblot, signal was quantified relative to 0h control (n=3 differentiations). Ordinary one-way ANOVA followed by Dunnett’s multiple comparisons test; for p-p38/p38: F(4,10) = 0.2660, P = 0.8932; peIF2a/eIF2a F(4,10)= 0.3824, P=0.8164. All comparisons n.s. Data are represented as mean ± SEM. **p)** Representative polysome profile of untreated hCO under basal conditions at day 25. Lysates were separated by ultracentrifugation through a 10–50% sucrose gradient, and absorbance was measured at 254 nm. Peaks corresponding to the 40S and 60S ribosomal subunits, 80S monosomes, and polysomes are indicated.

We next asked whether Emetine directly reduced nascent protein synthesis under the conditions that altered *EOMES*. We used the noncanonical amino acid β-ethynylserine (βES) to label newly synthesized polypeptides as a measure of translation output (Fig. 4j)^37^. Following βES incorporation, hCO treated with Vehicle or Emetine were dissociated, subjected to click chemistry, and analyzed by flow cytometry (Fig. 4k). βES labeling was consistently reduced following Emetine treatment across four independent differentiations, confirming that Emetine inhibits protein synthesis at a concentration of 250 nM across 6 days of treatment used to alter cortical cell-state progression.

Because perturbation of translation can activate cellular stress response pathways, we next examined the integrated stress response (ISR), ribotoxic stress response (RSR), and ribosome-associated quality control (RQC) pathways^38^ (Fig. 4l). hCO were exposed to Emetine for 1, 3, 6, or 24 hours as these pathways are engaged within minutes to a few hours^39,40^ and analyzed by immunoblotting for canonical markers of ISR and RSR (Fig. 4m) and RQC (Fig. 4n) activation. We detected no RPS3 ubiquitylation as an RQC marker during this time course, and quantification of ISR- and RSR-associated markers likewise revealed no significant activation across the time points examined (Fig. 4o). Thus, the early response to Emetine occurred without detectable activation of these canonical stress pathways. This is consistent with the fact that other translation elongation inhibitors, such as Cycloheximide (Fig. 4e) and Anisomycin (Fig. S5d) that typically induce these stress response pathways^38^ had no effect on *EOMES* expression.

To further examine the translational state of hCO during this developmental window, we performed polysome sucrose gradient fractionation under baseline conditions at day 25 (Fig. 4p) and on a positive control lysate (Fig. S5i) from a cell line known to robustly activate RQC^41^. The hCO profile showed small polysome fractions, indicating relatively low baseline ribosome occupancy in these fractions. This low abundance of densely loaded ribosomes may contribute to the absence of detectable acute RQC following Emetine treatment.

### Temporal characterization of translational and transcriptional responses to Emetine

Our previous analyses did not identify a canonical stress pathway that accounted for the effect of Emetine on neural progenitor progression. Since Notch signaling is a major regulator of cortical progenitor maintenance and neurogenic progression^42^, we first asked whether the response to Emetine could be achieved through modulation of this pathway. hCO were treated for 6 days with Emetine, the γ-secretase inhibitor DAPT, or combined Emetine and DAPT. DAPT increased *TBR1* expression, consistent with enhanced neuronal differentiation following Notch inhibition (Fig. S5j) as previously reported^43^. Emetine and DAPT produced an additive increase in *EOMES* expression (Fig. S5k), suggesting that the effect of Emetine on progenitor progression is not fully accounted for by Notch inhibition. To define the molecular changes associated with Emetine treatment over time, we performed bulk RNA sequencing of hCO after 3 h, 6 h, 24 h, 2 d, 4 d, and 6 d of treatment. In parallel, we profiled newly synthesized proteins after 3 and 6 h by labeling nascent polypeptides with βES, followed by biotin-azide click chemistry, tandem mass tag (TMT) labeling, and quantitative mass spectrometry (Fig. 5a). Independent assessment of βES incorporation confirmed reduced nascent protein synthesis at both 3h (Fig. S6a-c) and 6h (Fig. S6d-f) following Emetine treatment. Quantitative analysis of the nascent proteome revealed median Log_2_FC of –0.281 and –0.425 at 3 and 6h, respectively (Fig. 5b, Table S5 and Table S6), consistent with the global reduction in protein synthesis measured by βES labeling. However, certain proteins showed a more robust change in translation at 6h Emetine treatment (52/5014 identified proteins from Table S6, Log_2_FC < –0.7, adjusted P < 0.05). We searched for unique features in transcripts encoding the top inhibited proteins; however, we found no enrichment in transcript length, CDS length, 5′UTR or 3′UTR length, or in amino acid composition (Fig. S6g, h).

**Fig. 5:**
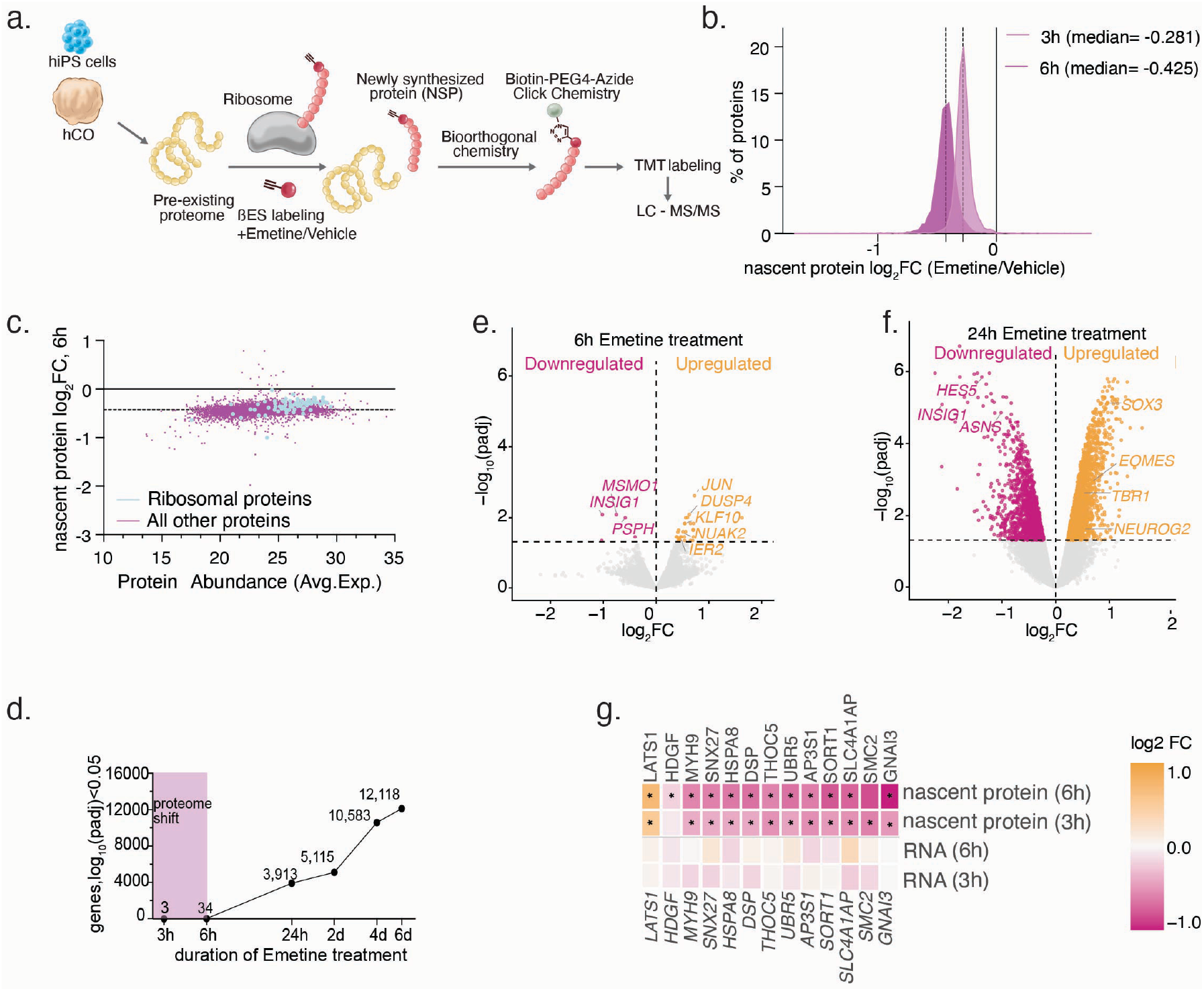
Temporal investigation of proteomic and transcriptomic changes in response to Emetine. **a)** Experimental design schematic for nascent proteome profiling using βES labeling. **b)** Histogram showing nascent proteome distribution across 3 h and 6 h timepoints where dotted lines indicate median Log_2_FC (-0.281 for 3h and -0.425 for 6h). **c)** Nascent protein Log_2_FC following Emetine treatment at 6h against protein abundance (AveExpr). Each dot represents a protein. Ribosomal proteins highlighted in blue (n = 79) are less affected than the remainder of the proteome represented in gray (n = 4,935). Median Log_2_FC −0.32 versus −0.425. Significance was assessed by comparing the observed difference in medians with a null distribution generated by randomly sampling 79 abundance-matched non-ribosomal proteins, over 5,000 permutations; no permutation reached the observed difference (one-sided permutation P = 0.0002). Dashed line indicates the median proteome-wide response (−0.425). **d)** Number of differentially expressed genes (FDR < 0.05) at each time point of Emetine treatment, from bulk RNA sequencing of hCO. Box highlighted in pink indicates timepoints measured by nascent proteome profiling. **e)** Volcano plot of transcriptional changes at 6h of Emetine treatment. Dotted line indicates FDR = 0.05. Of the 34 differentially expressed genes, several are immediate-early genes (*JUN, DUSP4, KLF10, NUAK2, IER2*) and metabolism-related genes (*INSIG1, MSMO1*). **f)** Volcano plot of transcriptional changes at 24h of Emetine treatment. Dotted line indicates FDR = 0.05. Neurogenic transcription factors (*EOMES, TBR1, NEUROG2*) are induced, while the Notch effector *HES5*, metabolism (*INSIG1*) and *ASNS* are reduced. **g)** Nascent protein synthesis (top) and transcript abundance (bottom) at 3h and 6h of Emetine treatment for proteins deviating reproducibly from the global translational response. Protein values are Log_2_FC, positive values indicate relative preservation of nascent synthesis, negative values greater suppression than the proteome. Asterisks denote FDR < 0.05 for the change in nascent synthesis relative to Vehicle.

Although the predominant response was a proteome-wide reduction in nascent protein synthesis, the magnitude of this response was not identical across proteins. At 6h, ribosomal proteins were relatively preserved compared with the overall proteome (median log_2_FC, -0.32 versus -0.425; Fig. 5c). We therefore asked when transcriptional changes emerged relative to this early translational response. Despite the pronounced change in nascent protein synthesis at 3 and 6h, bulk RNA sequencing identified only 3 and 34 differentially expressed genes, respectively, at FDR < 0.05. The transcriptional response subsequently expanded substantially, with 3,913 differentially expressed genes at 24 h, 5,115 at 2 d, 10,583 at 4 d, and 12,118 at 6d (Fig. 5d). Thus, widespread transcriptional remodeling emerged after the initial reduction in nascent protein synthesis.

The limited transcriptional response at 6h included immediate-early response genes such as *JUN* and *IER2*, together with a small number of regulatory and metabolic transcripts (Fig. 5e). By 24h, the transcriptional response had expanded to include genes associated with cortical progenitor and neuronal state transitions, including increased *EOMES, NEUROG2*, and *TBR1*, together with changes in genes such as *HES5* (Fig. 5f), a marker of neural progenitors. These results place the broad transcriptional response downstream in time from the initial perturbation of protein synthesis.

We finally examined whether individual proteins reproducibly deviated from the predominant proteome-wide response. A small subset of proteins showed concordant relative preservation or enhanced suppression at both 3 and 6h (Fig. 5g). Among these, LATS1 showed the most pronounced reproducible relative increase in nascent synthesis despite the global decrease in protein production, whereas *LATS1* transcript abundance remained unchanged at these early time points. Thus, Emetine produces a predominantly global reduction in nascent protein synthesis accompanied by a smaller reproducible set of protein-specific deviations, preceding the widespread transcriptional changes associated with altered cortical cell-state progression.

## Discussion

Here, we establish a scalable, semi-automated platform for arrayed perturbation of hCO and demonstrate its ability to uncover regulators of transient developmental cell states. Screening 1,280 small molecules for regulators of EOMES expression using a quantitative readout identified Emetine as the strongest hit, and this response was reproducible across multiple hiPS cell lines, macaque-derived organoids, and primary human fetal cortical tissue. Single cell profiling further showed that Emetine shifted cortical states away from dividing neural progenitors toward early neuronal populations. Additionally, the platform addresses a major limitation of organoid models: although they increasingly resolve human developmental cell states with high fidelity^1,2,44-46^, systematic perturbation at scale has remained challenging^13^.

Although Emetine is a well-established inhibitor of translation elongation^32^, its effect on cell state transitions was not a consequence of reduced protein synthesis in general, as only a subset of translation inhibitors increased *EOMES*. Emetine reduced nascent protein synthesis without detectable activation of the pathways associated with proteotoxicity. Nascent-proteome profiling revealed broad suppression of protein synthesis within 3-6 h, when comparatively few transcriptional changes were detectable. Widespread transcriptional remodeling emerged subsequently and included regulators associated with progenitor and early neuronal states. A small subset of proteins reproducibly deviated from the global translational response, including LATS1, a core regulator of the Hippo signaling pathway, which showed an increase in the βES-labeled protein fraction despite no corresponding early change in its transcript. This increased recovery could reflect changes in synthesis, altered degradation of newly synthesized LATS1, or both. Moreover, downregulation of LATS1, and the downstream Hippo signaling pathway, has been implicated in an increase in dividing neural progenitor populations in the developing mouse brain^47^; therefore, the Hippo pathway might also be relevant for human neural progenitor progression. Taken together, these findings place altered protein synthesis temporally upstream of the broader transcriptional and cell-state responses while leaving open which individual translational events contribute functionally to the phenotype.

Our study has several limitations. First, Emetine represents a pharmacological perturbation of elongation and there is little evidence for how the observed translational remodeling contributes to similar cell-state transitions under physiological conditions. Interestingly, translation and protein quality control are dynamically regulated during corticogenesis, including translational control within RG progenitors and neuronal fate regulators^48-53^. Determining whether endogenous changes in translation rate, ribosome composition, or the synthesis of specific proteins regulate progenitor-to-neuron transitions will require direct analysis under unperturbed developmental conditions in hCO^54^. In addition, our findings raise the possibility that translational control may contribute to modulating the rate of corticogenesis across species. Such studies remain technically challenging because hCO contain a diversity of cell types that must be isolated for cell type-specific analysis, typically requiring dissociation. This manipulation can itself alter rapidly regulated translational states and thereby confound measurements of immediate changes in protein synthesis. The relevance of translation in corticogenesis is further underscored by neurodevelopmental disorders caused by disruption of distinct components of the translational machinery^55^, including disorders caused by the genes *FMR1*^52,56,57^, *RPL10*^58^, *EIF2S3*^59^, *NEMF*^60^ and *DDX3X*^61^. These disorders point towards the sensitivity of neurodevelopment to perturbations in protein synthesis and raise the possibility that transient neural progenitor states are particularly vulnerable to fluctuations in translation rates^62^. Another limitation is that the present screen is focused on hCO and therefore does not establish whether similar regulatory principles operate across other regions. Importantly, however, the modular nature of this platform makes this limitation experimentally addressable and should enable its application to other regionally patterned organoids and assembloids.

Taken together, our study highlights the importance of translation in neural development and raises the question of whether modulating translation may have therapeutic relevance in certain neuropsychiatric disorders.

Looking forward, combining scalable perturbation with cell-type-resolved molecular analyses could help uncover the mechanisms governing human neural cell-state transitions.

Extending this framework across brain regions and developmental contexts may reveal both shared and context-specific regulatory principles, including selective cellular vulnerabilities.

## Acknowledgements

We thank all members of the Paşca lab for helpful discussions. We thank the Stanford Gene Vector and Virus Core for amplification of the *EOMES* lentivirus vector. *EOMES* LV vector was a gift from Feng Zhang. We thank the Genomics Service Center for Bioanalyzer analysis for single-cell RNA-seq experiments and the Stanford High-Throughput Nucleus for help with acoustic dispensing and access to LOPAC1280 library and the Stanford FACS core for help with FACS experiments and CZ BioHub for sequencing the scRNA-Seq libraries.

## Author contributions

MA, SPP conceived the project and designed the experiments. MA developed the high-throughput organoid cultures and characterized them, established the screening platform and performed the screens, differentiated hiPS cell lines and dissociated hCO, prepped the scRNA-Seq libraries and performed FACS experiments for bulk RNA-Seq, DAPT and Emetine experiments, embedding and cryosectioning of organoids. MA and TL performed βES-labeled nascent proteomics experiments, western blotting. TL performed click chemistry on βES-labeled organoids and downstream FACS analysis and polysome profiling and ISR, RSR, RQC western blots. DH helped perform the screens and performed FACS analysis for bulk RNA-Seq experiments and TBR2 western blot. DH and MO performed the immunostainings, imaging and quantification. MO and ML performed live imaging. KI performed mqCO experiments and DNA extraction for SNP array analysis of ER. VK assisted in scanner imaging of organoids, maintenance of organoids, qPCR readouts. DW assisted with mqCO differentiations. MF analyzed bulk RNA-Seq and scRNA-Seq data. SY differentiated hCO, MA, CD and RJL performed primary tissue experiments. ES derived the mqiPS cell lines, AB ran the mass spectrometry experiment after sample prep. IL analyzed 6h nascent proteomics data. KK supervised MF for scRNA-Seq analysis, ATC supervised ES, SP supervised AB, An.B supervised DH, MB supervised aspects of the study related to translation inhibition and IL and TL. MA and SPP wrote the manuscript with input from all authors. SPP supervised all aspects of this study.

## Competing Interest Statement

Stanford University holds patents for the generation of neural organoids (listing S.P.P. as an inventor).

## Funding

This work was supported by an NIH grant (R01MH107800), the Stanford Big Idea Project on Brain Organogenesis (Wu Tsai Neurosciences Institute) (to S.P.P.), the Kwan Funds (to S.P.P.), the Senkut Funds (to S.P.P.), an SNSF Postdoc.Mobility grant (222016 to M.A.), an EMBO Postdoctoral fellowship ALTF 1045-2022 and SNSF Postdoc.Mobility Grant 230454 (to D.H.). Pew Innovations Award (to M.B.). The funding for generating the rhesus cell lines were from the NIH (R01HD098278 and P01HD075795) to A.T.C. S.P.P. is a CZ BioHub Investigator and a CZI Ben Barres Investigator.

## List of Supplementary Materials

Materials and Methods

Tables S1-S6

Movie S1

## Materials and Methods

### hiPS cell culture and neural differentiation

Human and macaque iPS cell lines (KOLF2.1J, 0524-1, riPSC05, riPSC89, 1208-2, 8119-1, 1205-4) were maintained and passaged as previously described^15^. The EOMES–NLS-tdTomato reporter (ER) iPS cell line was generated by Bayspair Inc. (Mountain View, CA, USA)^63^. *EOMES* alleles were modified by an in-frame insertion of a T2A–NLS–tdTomato cassette immediately upstream of the endogenous EOMES stop codon, resulting in a biallelic EOMES reporter line. Correct targeting was confirmed by genomic PCR using a primer pair located outside the 5′ and 3′ homology arms. The size of the resulting PCR product was consistent with the expected targeted allele, and the edited locus was further verified by Sanger sequencing. Cells were tested regularly to ensure they were maintained Mycoplasma-free. Approval for the derivation and/or use of these cell lines was obtained through the Stanford Institutional Review Board. Genome integrity of the ER cell line was assessed at the Children’s Hospital of Philadelphia using the Illumina genome-wide GSAMD-24 v2.0 SNP microarray. Bulk hCO cultures were generated and maintained using previously established protocols^15^. mqCO were generated and maintained identical to the hCO protocol.

For high throughput organoid cultures, 30,000 cells/well were seeded in low attachment U-Bottom 96-well plates (7707, Corning) in 100 µL of Essential 8 medium supplemented with ROCK inhibitor Y27632 (10 µM, Selleckchem, S1049) and this was designated as D0 for differentiation and plates were centrifuged at 100x g for 3 minutes. In all subsequent media changes, a semi-automated plate washer dispenser (Biotek EL406) was used to aspirate from and dispense to 96-well plates. On d1, media change with Essential 6 media (Thermo Fisher, A1516401) supplemented with 2.5 *µ*M DM (dorsomorphin, Sigma Aldrich, P5499) and 10 *µ*M SB (SB-431542, R&D Systems, 1614) was done twice to remove all growth factors remaining from the Essential 8 media in the wells. All other guidance molecules and concentrations and durations for the molecules were kept identical to the previously published protocol^15^ with the exception that from d7 onwards media was changed every other day for the duration of the cultures. Imaging of plates for quantification of growth of organoids was done using an Epson Perfection V850 Pro Scanner and analyzed using ImageJ 1.53K software.

### Primary tissue culture

Human tissue was obtained under a protocol approved by the Research Compliance Office at Stanford University. GW17 cortical tissue was delivered on ice and processed immediately after arrival. Tissue was sectioned at 500 µm using a Leica VT1200S vibratome and transferred onto cell-culture membrane inserts (0.4-µm pore size; Falcon) in 6-well plates (Corning). Slices were maintained in the same neural medium used for hCO cultures, supplemented with 1% Antibiotic-Antimycotic (Thermo Fisher Scientific) during the first 6 days in culture.

### LOPAC1280 and translation inhibitor library reformatting

LOPAC1280 library (Sigma Aldrich) was acquired from the Stanford High-Throughput Screening Core Facility at the Nucleus at Sarafan/Chem-H building. The library was reformatted into intermediate 96-well plates (Greiner BioOne PP, 650201) using acoustic dispensing (Beckman Coulter, Echo655 liquid handler) and stored at –20°C until use. For the secondary screen, we reformatted Proglumide (Sigma, M006) and substituted 6-Fluo-ronorepinephrine with Norepinephrine (Sigma, A0937) as this product was discontinued. Translation inhibitors (Emetine; E2375, Sigma, Anisomycin; S7409, Selleckchem, Cryptopleurine; HY-121255, MedChem Express, Cycloheximide; S7418 Selleckchem, Homoharringtonine; HY-14944, Med-Chem Express, 4EGI-1; HY19831 MedChem Express, Didemnin B; HY105055 MedChem Express, Puromycin; S7417 Selleckchem, INK128) were resuspended and transferred to either Eppendorf tubes (Fisherbrand; 05-408-129) as aliquots or ECHO-compatible 384-well Polypropylene intermittent reformatting plates (00114615, Beckman Coulter) to be reformatted into intermediate 96-well plates (Greiner BioOne PP, 650201) using acoustic dispensing (Beckman Coulter, Echo655 liquid handler) and stored at -80°C until use.

### Semi-Automated compound screening in cortical organoids

hCO were seeded and cultured as described above. On day 25, 27 and 29 of culture, reformatted library plates were thawed, centrifuged at 100x g for 1 min and diluted in media using a Viaflo 96 liquid handling robot (Zizers, Switzerland) to make 10 *µ*M and 500 nM stocks for the high and low concentrations, respectively. Following medium change on the hCO plates, compounds were added from the reformatted compound stock plates using a Viaflo liquid handling robot. On day 31 of culture, hCO were washed 2x with DPBS (14190144, Thermo Fisher Scientific) using the BiotekEL406 and were lysed with the Cells-to-Ct 1-Step Taqman lysis buffer (A25603, Thermo Fisher Scientific) using Viaflo. Lysis was stopped with the Stop reagent from the same kit (A25603, Thermo Fisher Scientific) and plates were centrifuged at 100x g for 2 min. These plates were denoted as stock lysate plates and an 11 *µ*L aliquot of these plates were transferred into ECHO-acoustic dispenser compatible low-dead volume (LDV) plates (001-16128, Beckman Coulter) and both sets of plates were stored at –80°C until further use.

### *μ*qPCR and viability readout for screening in cortical organoids

LDV plates were thawed and centrifuged at 100x g for 2 min prior to ECHO-acoustic dispenser reformatting. Taqman Fast Virus 1-Step Master Mix (4444436, Thermo Fisher Scientific) was diluted in water and a total of 3.6 *µ*L was dispensed/well into destination plates (MicroAmp Optical 384-well qPCR Reaction Plate, 4309849, Thermo Fisher Scientific) using a Certus Flex multidrop dispenser (Gwatt, Switzerland). FAM-MGB Taqman Assay Probes (human *EOMES, GAPDH, FOXG1*) were aliquoted into ECHO-compatible 384-well Polypropylene intermittent reformatting plates (00114615, Beckman Coulter) and a total of 200 *n*L/assay was finally reformatted using an acoustic dispenser into the MicroAmp qPCR plates. Lysates were reformatted from the LDV plates to the MicroAmp qPCR plates using an acoustic dispenser at a volume of 200 *n*L/assay to make up a final reaction volume of 4 *µ*L/qPCR assay. Plates were read out using QuantStudio™ 5/6/7 Real-Time PCR Systems (Thermo Fisher Scientific) using the following cycling conditions: hold Stage: 50°C (02:00), 95°C (10:00) and PCR Stage (40-cycles): 95°C (00:15), 60°C (1:00). Data were analyzed following the ddCq method^64^. Plate 42 was omitted from further analysis as it did not pass quality control and wells with unusually high values for *GAPDH*, indicating plate transfer problems were also disregarded from further analysis. Details for probes used can be found in Table S2. Viability was assessed using 3D-Glo kit (G9683, Promega) according to instructions from the manufacturer. In brief, the viability reagent and plates containing organoids were equilibrated to RT and 125 *µ*L of media was discarded from the wells, 100 *µ*L viability reagent was added and the plate was put on plate shaker for 5 minutes at 700 rpm (Eppendorf Thermo Mixer C) and 25 min further incubated at RT to stabilize luminescence signal. A total of 200 *µ*L of supernatant was transferred to opaque (6005660, Perkin Elmer 96-well CulturPlate) plates for recording signal. Luminescence was recorded on a Varioskan LUX plate reader (Thermo Fisher Scientific).

### Two-step conventional qPCR with SYBR^™^ green

SYBR-green based qRT-PCR was performed as previously described^65^. Up to three hCO were pooled in the same tube and were considered one sample. RNA was extracted using RNeasy Mini Kit (74106, Qiagen). cDNA synthesis was done using SuperScript^™^ III First-Strand Synthesis SuperMix for qRT-PCR (Thermo Fisher Scientific, 11752250) with DNase I treatment step. qPCR was done using SYBR^™^ Green PCR Master Mix (Thermo Fisher Scientific, 4312704) on a QuantStudio™ 6 Real-Time PCR System (Thermo Fisher Scientific). Details for primers used can be found in Table S3.

### Cryopreservation, immunocytochemistry (ICC) and imaging

hCO were fixed in 4% paraformaldehyde (PFA) in DPBS (15710, Electron Microscopy Services) for 30 minutes at room temperature (RT). Following fixation organoids were washed twice with DPBS and transferred to a solution containing 30% sucrose (S9378, Sigma Aldrich) and left at 4°C until they sank to the bottom for cryopreservation. hCO were then embedded in a 1:1 (vol/vol) mix of optimal cutting temperature compound (OCT, 23-730-571, Fisher Scientific) and 30% sucrose. Blocks were frozen on dry ice, and 12-15 µm thick sections were cut onto Superfrost Plus microscope slides (VWR, 48311-703) using a Leica Cryostat. For immunocytochemistry, sections were blocked for 1h at RT using 1% BSA (A9418, Sigma Aldrich), 0.1% Triton X-100 (AC327372500, Thermo Fisher Scientific) and 0.02% Sodium Azide (RTC000068, Sigma Aldrich) diluted in DPBS. Slides were incubated overnight at 4°C with primary antibodies in blocking solution. The next day, sections were washed with blocking solution and further incubated with Alexa Fluor coupled secondary antibodies (1:250 dilution in blocking solution) for 1h at RT and washed three times with DPBS. Nuclei were visualized with DAPI (Sigma Aldrich) and slides were mounted using Aqua Poly Mount (18606, Polysciences). Antibodies used in this study can be found in Table S4. Imaging of sections was performed on a Zeiss Axioscope widefield, EVOS M7000 slide scanner, Leica Stellaris 5, Leica SP8, or Zeiss LSM9 Series confocal imager. For live imaging organoids were transferred to 96-well imaging plates (Ibidi µ-Plate 96 Well imaging plate ibiTreat, 50-305-829, Fisher Scientific) and samples were imaged under controlled environmental conditions (37°C, 5% CO_2_) using a 10x objective on a Leica Stellaris confocal imager.

### ICC Image analysis

Image analysis was done with Fiji (Java 21.0.7). For co-stain analysis, tdTomato and TBR2 channels were thresholded and thresholded images were converted to binary masks. The tdTomato mask was subjected to watershed segmentation to separate adjacent tdTomato-positive objects. tdTomato-positive particles were then identified using the Analyze Particles function with a minimum particle size of 20 pixels and no circularity restriction (0–1), and individual particle regions of interest (ROIs) were added to the ROI Manager. TBR2 positivity was quantified within each tdTomato-defined ROI using the corresponding binary TBR2 mask. For each tdTomato-positive particle, the fraction of the ROI occupied by TBR2-positive pixels was calculated from the mean intensity of the binary TBR2 mask, and the overlapping area between the tdTomato and TBR2 masks was calculated as the tdTomato ROI area multiplied by the TBR2-positive fraction. tdTomato-positive particles with ≥20% overlap with the TBR2 mask were classified as TBR2-positive. The percentage of tdTomato-positive particles that were TBR2-positive was calculated for each image. All other quantifications were performed in a similar fashion, by thresholding single channel images and assessing either positive particles using the Analyze Particle function or area with the measuring function and dividing by the same metric in the DAPI channel to normalize to total cell amounts. All images within an experiment were analyzed using the same segmentation and classification parameters and blindly.

### Dissociation of hCO

Four to eight hCO that were d31 and d50 of neural differentiation for single cell experiments and d31of neural differentiation for bulk RNA-Seq experiments were transferred into low attachment 6-well plates (Corning, 3471, Fisher Scientific) and digested using papain solution (30 U ml^−1^ papain (LS003127, Worthington Biochemical), 10x Earle’s balanced salt solution (Sigma-Aldrich, E7510), 0.46% D(+)-glucose, 0.5 mM ethylenediaminetetraacetic acid (EDTA), 25 mM NaHCO_3_, 125 U ml−1 Deoxyribonuclease I (Worthington Biochemical, LS002007) and 6.1 mM L-cysteine (Sigma-Aldrich, C7880) for 45 min at 37°C as previously described^15^. Following papain digestion, organoids were washed twice with Enzyme inhibitor solution (2% trypsin inhibitor, Worthington Biochemical, LS00308, 10x Earle’s balanced salt solution, 0.46% D(+)-glucose, 25 mM NaHCO_3_) and centrifuged at 100x g for 30s in between washes. After washes, organoids were gently triturated with serial pipetting four to five times to achieve a single cell suspension in a low-ovo solution. Single-cell suspension of hCO was subjected to gradient based centrifugation in a highovo solution to achieve pelleting following centrifugation at 200x g for 5 min.

### scRNA-Seq and bulk RNA-Seq

Pelleted single-cell suspension was resuspended in 0.04% BSA (A1595, Sigma Aldrich) in DPBS and filtered through a Flowmi pipette tip filter (H13680-0040, Bel-Art). Cells were then counted using a Countess^™^ Automated cell Counter (Thermo Fisher Scientific) using Trypan blue staining (15250061, Thermo Fisher Scientific). Cells were loaded onto a Chromium Single Cell 3′chip (Chromium Next GEM Chip G Single Cell Kit, 10X Genomics, PN-1000127). cDNA libraries were generated with a Chromium Next GEM Single Cell 3′ GEM, Library & Gel Bead Kit v3.1 (10X Genomics, PN-1000128 and PN1000269), following manufacturer’s instructions. cDNA quality and quantity were assessed using the Agilent Bioanalyzer High Sensitivity DNA Assay through the Stanford Genomics Core Facility. Barcoding of samples was done using the dual index kit TT Set A (1000215, 10X Genomics). Libraries were sequenced at the CZ BioHub in San Francisco using an Illumina NovaSeq X Sequencer.

For bulk RNA-Sequencing following FACS isolation, cells were dissociated into a single cell suspension as described above. Following dissociation, cells were sorted into tdTomato^+^ and tdTomato^-^ populations at the Stanford Shared FACS facility using FACSAria II, FACSAria Fusion. Analysis was done with FlowJo (v.10.8.1) software. Bulk RNA-Sequencing sample prep and sequencing was performed by Novogene Inc. In brief, RNA was extracted from pelleted cells and Poly-A enriched mRNA was sequenced on the Illumina NovaSeq X Plus Series sequencer (PE150) with 6G raw data per sample. For time-resolved bulk RNA-Seq, samples were preserved in RNA/DNA shield (R1100-50, Zymo Research) and sent to Plasmidsaurus Inc. to be analyzed.

### scRNA-seq data analysis

#### Preprocessing, quality filtering and sample integration

Raw reads were aligned to the GRCh38-2024-A reference genome, and UMI (unique molecular identifier) counting was performed using the Cell Ranger (10X Genomics version 9.0) pipeline with default parameters. Downstream analyses were performed using R (version 4.4.2) and Seurat (version 5.2.1)^66^. For each individual sample we first removed cells with high mitochondrial content (>10%) and cells with less than 500 genes. We then applied the DoubletFinder (version 2.0.4)^67^ approach to doublet removal. Raw gene count matrices were normalized by regularized negative binomial regression using the *SCTransform* function (vst.flavor=“v2”), which also identified the top 3,000 highly variable genes using default parameters. Further filtering rounds were performed within each sample of clusters of putative low-quality cells based on outlier low unique gene counts (median below the 10th percentile). To integrate the samples within each corresponding cohort (D31 with differentiation 1 and 2, and D50), we used Seurat *SelectIntegrationFeatures* function considering *nfeatures* = 3,000. The integration anchors were determined by sample library using parameters *normalization*.*method* = “SCT”, *reduction* = “rpca” and the 3,000 selected features. We then considered the *IntegrateData* function using *normalization*.*method* = “SCT” and 50 dimensions as parameters. Another round of putative low-quality filtering, based on clusters with outlier low gene counts (median below the 10th percentile), was performed on the integrated Seurat object. The clusters were determined based on *FindNeighbors* (using 50 dimensions as parameter) and *FindClusters* functions with a resolution of 3. Analogous preprocessing and filtering were performed on the D31 and D50 sample cohorts. Uniform Manifold Approximation and Projection (UMAP)^68^ was used for visualizing cell types and clusters with representative markers.

#### Cell clustering and annotation

We then used the Seurat object to define an initial cluster considering Seurat’s *FindNeighbors* (using 50 dimensions as parameter) and *FindClusters* function with a resolution of 2. We then performed a Differential Gene Expression analysis using *FindAllMarkers* function with Wilcoxon test, *min*.*pct* = 0.10 and *logfc*.*threshold* = 0.25 parameters. We identified and categorized major cell classes through the presence of the following marker gene signatures. Dividing progenitor clusters displayed expression of mitotic transcripts *MKI67* and *TOP2A* and/or *SOX2*; Intermediate progenitor cells (IPC) expressed *EOMES, NEUROG1;* Forebrain GABAergic neurons expressed GABA processing enzymes *GAD1, GAD2*; Cortical hem and Choroid expressed *LMX1A, WNT5A, TTR*; other non-cortical neurons expressed *TFAP2A, TFAP2B;* hindbrain glutamatergic neurons expressed *ATOH1;* Early cortical glutamatergic neurons expressed *FOXG1, SLC17A6, SLC17A7, TBR1;* Cajal-Retzius cells expressed *TP73, RELN*; Radial glia expressed *FOXG1, HES1, SOX2;* Midbrain progenitors expressed *IRX1, IRX2* and Other progenitors; non-*FOXG1* with *SOX2*.

#### Differential gene expression analysis

Differential gene expression analysis between Emetine and Vehicle-treated hCO was performed using a pseudobulk approach across samples. Seurat *AggregateExpression* function was used to sum counts for each broad cell type annotation. We used DESeq2 (version 1.46.0) to normalize and perform differential expression taking into account cell lines and differentiation as covariates for the d31 cohort. Genes with counts lower than 10 were filtered out. GO term enrichment analysis was performed with clusterProfiler R package (version 4.14.6) using gene lists with adjusted P-value <0.05 and absolute log2 fold-change > 0.5.

#### Pseudotime inference

To assess the differences in pseudotime associated with Emetine treatment we performed trajectory inference using Slingshot (version 2.14.0), tradeSeq (version 1.20.0) and condiments (version 1.14.0) R packages. We first subset cells of interest and recluster using Seurat *FindNeighbors, FindClusters* and *RunUMAP* procedures on the integrated assay. Slingshot was applied over the UMAP reduced dimension considering dividing progenitor cell as a starting cluster. To test for differential pseudotime progression between conditions (Fig. 3i and Fig. S4j), we emulate the condiments *progressionTest* function to shuffle condition labels across sample-level permutations instead of individual cells. The permutation test takes into consideration the cell lines and differentiations as pseudo-bulk samples. We first calculate weighted mean pseudotime per sample, next we permute ‘condition’ labels across the pseudo-bulk samples (2000 permutations) and test whether the observed mean pseudotime is different from the empirical null distribution.

#### Cell compositional analysis test

To assess the differences in cell composition (Fig. 3e) we developed a Linear Mixed Model (LMM) using lme4 (version 2.0.1) and lmerTest (version 3.2.1) R packages. Proportional counts associated with each sample and cell type were arcsine-square-root transformed. We then fit the LMM taking into consideration cell line and differentiation batch as random effects and condition as a fixed effect. The estimation was based on Maximum Likelihood (ML). We discarded groups with cells that lacked observations in both conditions. To determine the direction of enriched or depleted cells in Emetine treatment we considered the Log_2_FC of the estimated marginal means using emmeans (2.0.3) R package. A pseudocount 0.001 was added to the arcsine transformed proportions.

### Bulk RNA-seq data analysis

#### Preprocessing and quality filtering and DEG analysis

Data quality assessment and sequence read primer trimming were performed by FastQC (version 0.11.9) and Cutadapt (version 4.9), respectively. Trim_galore version 0.6.7 with *--phred33* parameter was considered for low-quality reads removal. The data was then aligned to a human reference genome build (*Homo sapiens* GRCh38). STAR (version 2.7.10b) aligner index was created using *STAR_index --genomeFastaFiles Homo_sapiens*.*GRCh38*.*dna*.*primary_assembly*.*fa --sjdbGTFfile gencode*.*v44*.*primary_assembly*.*annotation*.*gtf* and a gene-level read count matrix was generated using featureCounts subread (version 2.1.1)^69^ commands. Differential gene expression and functional enrichment analysis were performed using R (version 4.4.2), Bioconductor (version 3.20). Differential expression was done with DESeq2 (version 1.46.0), low-expressed genes (counts ≤ 5 and at least 50% of the samples) were filtered out and the differential expression analysis performed using *DESeq* function. Differentially expressed genes were selected using an absolute log2 fold change of ≥ 1 and an adjusted P value of ≤ 0.05.

#### Time course single-end RNA-Seq processing

Quality of the fastq files was assessed using FastQC (version 0.12.1). Reads were then quality filtered using fastp (version 0.24.0) with poly-X tail trimming, 3’ quality-based tail trimming, a minimum Phred quality score of 15, and a minimum length requirement of 50 bp. Quality-filtered reads were aligned to the reference genome using STAR aligner (version 2.7.11) with non-canonical splice junction removal and output of unmapped reads, followed by coordinate sorting using samtools (version 1.22.1). PCR and optical duplicates were removed using UMI-based deduplication with UMIcollapse (version 1.1.0). Alignment quality metrics, strand specificity, and read distribution across genomic features were assessed using RSeQC (version 5.0.4) and Qualimap (version 2.3), with results aggregated into a comprehensive quality control report using MultiQC (version 1.32). Gene-level expression quantification was performed using featureCounts (subread package version 2.1.1) with strand-specific counting, multi-mapping read fractional assignment, exons and three prime UTR as the feature identifiers, and grouped by gene_id. Final gene counts were annotated with gene biotype and other metadata extracted from the reference GTF file. Differential expression was done with edgeR v4.0.16 using standard practice including filtering for low-expressed genes with edgeR::filterByExpr with default values.

### β-Ethynylserine (βES) labeling, flow cytometry

To measure translation rates, βES (L-β-Ethynylserine, HY-164745, MedChem Express) was used at a final concentration of 2 mM to label nascent peptides. Three to four hCO per condition were dissociated as described above. Following dissociation, cells were pelleted, fixed with 4% PFA in FACS tubes for 15 min on ice subsequently washed and resuspended in 0.04% BSA in DPBS. Cells were centrifuged at 4°C at 1000x g for 5 min and pellets were resuspended in 500 uL of 0.1% freshly made Saponin (w/v) in PBS for permeabilization and incubated for 20 min on ice in the dark. 1.5x Click reaction without sodium ascorbate was prepared as follows: 187.5 µM CuSO_4_, 375 µM BTTAA, and 3.75 µM CALFLUOR 488 azide in PBS and 15 mM Sodium Ascorbate (3x) in PBS. CuSO_4_ and BTTAA should be prepared at least 5 min prior to dilution in PBS to allow chelation. Following permeabilization, samples were spun down at 1,400x g for 10 mins at 4°C and supernatant was aspirated following centrifugation. Click reaction was performed at RT by resuspending the pellet in 800 µL of the 1.5x click reaction as described above. Following resuspension of pellets in 1.5x click reaction, 400 µL of 15 mM sodium ascorbate was added to each sample and incubated for 1h in the dark at RT. Samples were centrifuged at 1,400x g at 4°C and click reaction was aspirated and pellet was suspended in 100 µL of FACS Buffer (PBS, 3% FBS, 2 mM EDTA) and samples were directly subjected to flow cytometry using the Novocyte Penteon (Agilent Technologies) system. Data were analyzed in FlowJo v10.10.0. Samples were gated on singlets (FSC-A vs. FSC-H and SSC-A vs. SSC-H) and down sampled to 14,000 cells using the DownSample v3.3.1 plugin. The median fluorescence intensity (MFI) of the entire single cell population was used to represent global protein synthesis. Within each independent experiment, background was subtracted using the MFI of a “no βES” control that underwent the click reaction.

### Western blotting for TBR2

hCO were harvested, pooled and frozen on dry ice until further use. Lysis was performed with 50 mM Tris–HCl, pH 8, 150 mM NaCl, 0.5% sodium deoxycholate, and 0.5% Triton-X 100, 0.1% SDS and a protease inhibitor (11836170001, Roche) was supplemented into the buffer. Further lysis was achieved via probe sonication (3 cycles of 8 pulses of 1 sec on/ 1 sec off at 35% amplitude). Samples were incubated on ice for 10 min followed by centrifugation at 10,000x g for 10 min. Protein concentrations were measured with a bicinchoninic acid assay (23225, Thermo Fisher Scientific). Samples were loaded on a 4–12% gradient gel (NP0323BOXA, Thermo Fisher Scientific). A semi-dry transfer method was used to blot the membrane (IB21001, Thermo Fisher Scientific) onto a PVDF membrane (IB24001, Thermo Fisher Scientific). Membrane was blocked with 5% BSA in 1X PBST (1706404XTU, Bio-Rad) for 30 min. Following blocking, primary antibodies were incubated overnight at 4°C with a 1:1000 dilution in 1% blocking buffer in 1X PBST (IBB-690, Boston BioProducts) and washed the next day before incubation with the secondary antibody. Fluorescent detection of GAPDH was done using a secondary antibody coupled to a near IR680 dye (Licor). Secondary antibody incubation was done at RT for 1h with a dilution of 1:10000 in 1% blocking buffer in PBST. Membranes were developed using an Odyssey Fc (Licor) imaging system and SuperSignal^™^ Femto/Pico PLUS chemiluminescent western substrate (34095 and 34580 respectively, Thermo Fisher Scientific). Antibodies can be found in Table S4.

### Time course of Emetine treatment of hCO with Sucrose Cushion for ISR/RSR/RQC

A total of 32 hCO following Emetine treatment at d25 of neural differentiation were pooled per condition, lysed in a cold, hypotonic polysome lysis buffer without detergents (20 mM Tris, pH 7.5, 5 mM magnesium chloride (MgCl_2_), 1 mM DTT, 10 mM potassium chloride (KCl), 5 mM Putrescine, 100 µg/mL cycloheximide, 200 U/mL SUPERase·In™ RNase Inhibitor, 1× mini complete protease inhibitor, 1x PhosSTOP, in nuclease-free water) and incubated for 5 min on ice. Next, samples were lysed further in a detergent and DNase containing lysis buffer (0.5% Triton X-100, 0.5% sodium deoxycholate, 20 U/mL TURBO DNase at 4°C) for 30 min with rotation end on end. Lysates were clarified through centrifugation at 1000x g for 5 min at 4°C. Supernatant was transferred to a new tube and the centrifugation step was repeated once more to further enable clarification of the lysates. For sucrose cushion, 180 µL of the clarified supernatant was loaded onto 700 µL 1M sucrose cushion buffer (1 M sucrose, 20 mM Tris, pH 7.5, 15 mM magnesium chloride (MgCl_2_), 1 mM DTT, 100 µg/mL cycloheximide, 200 U/mL SUPERase·In™ RNase Inhibitor) and centrifuged at 100,000 rpm at 4°C using a Beckman TLA120.2 rotor for 1h. The rest of the lysates were stored at -80 °C. The input lysate was used subsequently for phospho-eIF2α/ total eIF2α and phospho-p38/ total p38 and cushion was used for RPS3 western blotting. As positive control for RQC 200 nM Anisomycin treated HEK293T cell lysate was used and for ISR/RSR, lysate from A549 treated with 0.1% DMSO or 40µM Anisomycin was used as a negative and positive control, respectively. *Western Blot for ISR/RSR/RQC*

Protein concentrations were measured with a bicinchoninic acid assay (23225, Thermo Fisher Scientific). Samples were loaded on a 4–15% stain-free gradient gel (5678084, Bio-Rad) for phospho-eIF2α/ total eIF2α and phospho-p38/ total p38 or a hand-cast 0.5 mm thick, 12% SDS-PAGE made from 30% Acrylamide/Bis Solution, 29:1 (1610156, Bio-Rad). A semi-dry transfer method was used to blot the membrane (Trans-Blot Turbo Transfer System, Bio-Rad) onto a 0.2 µm PVDF membrane (1620177, Bio-Rad) using the “mixed MW” setting for the 4-15% stain-free gradient gel or “low MW” setting for the handcast 12% gel. Membrane was blocked with AdvanBlock™-Chemi (R-03726-E10, Advansta) for 60 min at room temperature. Following blocking and washing with 1x PBST, primary antibodies were incubated overnight at 4°C with a 1:1000 dilution in AdvanBlock™-Chemi and washed the next day before incubation with the anti-mouse (NA931-1ML, Cytiva) or anti-rabbit (NA934-1ML, Cytiva) HRP conjugated secondary antibody. Secondary antibody incubation was done at RT for 1h with a dilution of 1:5000 for RPS3, 1:10000 for P-eIF2α and P-p38, 1:15000 for eIF2α and p38, 1:20000 for β-actin in AdvanBlock™-Chemi. Following washing with 1x PBST, Membranes were developed using Clarity Western ECL Substrate (1705061, Bio-Rad) and SuperSignal^™^ Femto PLUS chemiluminescent western substrate (34095, Thermo Fisher Scientific). Antibodies can be found in Table S4.

### 10-50% Sucrose Gradient for Polysome Profiling

A total of 96 hCO at day 25 of neural differentiation per condition were collected in a conical tube and medium was removed. Polysome stimulation was done by adding 10 mL fresh culture media to hCO and incubated for 20 minutes at 37°C. hCO were treated with 100 µg/mL Cycloheximide for 3 mins at 37°C and media was aspirated in full and hCO were immediately resuspended in 400 µL of freshly prepared cold hypotonic polysome lysis buffer without detergents (20 mM Tris, pH 7.5, 5 mM magnesium chloride (MgCl2), 1 mM DTT, 10 mM potassium chloride (KCl), 5 mM Putrescine, 100 µg/mL cycloheximide, 200 U/mL SUPERase·In™ RNase Inhibitor, 1× mini complete protease inhibitor, 1x PhosSTOP, in nuclease-free water) and incubated for 5 min on ice. Next, samples were further lysed in a detergent and DNase containing lysis buffer (0.5% Triton X-100, 0.5% sodium deoxycholate, 20 U/mL TURBO DNase at 4°C) for 30 min with rotation end on end. Lysates were clarified by first spinning down at 1000x g for 5 min at 4°C. Supernatant was transferred to a new tube and lysates were further centrifuged at 21,300x g for 15 min and supernatant was transferred to a new tube. 10-50% sucrose gradient fractionation: 300 µL of lysate of hCO (corresponding to 46 µg of RNA, following quantification by QUBIT RNA HS assay (Qiagen), and 150 µg of RNA from the A549 positive control lysate) was layered onto a freshly prepared 10-50% (w/v) sucrose gradients (20 mM Tris pH 7.5, 150 mM NaCl, 15 mM MgCl2, 1 mM DTT, 100 µg/mL Cycloheximide, made on a Biocomp Model 108 Gradient Master using the preprogrammed long cap 10-50% sucrose gradient setting). Gradients were centrifuged in a Beckman SW41 rotor at 40,000 rpm for 2.5h at 4 °C in a Beckman L8–80M ultracentrifuge. Following centrifugation, gradients were fractionated into 16 fractions using a Biocomp Piston Gradient Fractionator with continuous A260 monitoring (Flow Cell software v2.50).

### Time course of Emetine treatment with βES coupled pulldown Mass Spectrometry for nascent proteome profiling

βES labeling was performed as described above for 3h and 6h during Emetine treatment. Longer βES labeling was toxic, therefore we could not pursue a longer labeling period. 48 hCO at d25 of neural differentiation per condition were pooled in a conical tube and medium was aspirated.

#### CuAAC click labeling with Biotin-PEG_4_-azide

Cell pellets were resuspended in 500 µL cold DPBS lysis buffer containing a 1× protease/phosphatase inhibitor cocktail (Thermo Fisher, 78443) and further lysed via probe sonication (8 pulses of 1 sec on/ 1 sec off at 35% amplitude) while taking precautions to prevent foaming. Protein concentrations were determined using a bicinchoninic acid (BCA) protein assay (Thermo Fisher, 23225) using BSA standards and normalized. For click chemistry labeling, a freshly prepared master reaction mixture containing 30 µL of 1.7 mM TBTA in DMSO-tBuOH (1:4 v/v), 10 µL of 50 mM CuSO_4_ in water, 5 µL of10mM Biotin-PEG_4_-azide in DMSO (ChemPep Inc., 271606), and 10 µL of 50 mM TCEP freshly prepared in water was added to each 500 µL proteome sample resulting in a final concentration of: 91 µM TBTA, 0.9 mM CuSO_4_, 91 µM Biotin-PEG4-azide, and 0.9 mM TCEP. The click reaction was allowed to proceed for 1h in the dark at 37°C with constant agitation. 2 mL of cold methanol was added to each click reaction of each sample and samples were incubated overnight at −20°C to precipitate proteins.

#### Quantification of protein synthesis inhibition via western blotting

For each sample in the βES mass spectrometry, 2x reactions with and without biotin-PEG4-azide click labeling were prepared. At 1/10^th^ scale, 5.5 µL click master mix (described above) or 5.5 µL of DPBS, respectively was added to 50 µL of 2 mg/mL protein. Click reaction was let to react for 1 hr at 37°C using a Bio-Rad T1000 thermocycler. Reaction was quenched using 19 µL 4x SDS loading buffer and loaded 10 µL onto a 4-15% Criterion TGX Stain-free Protein Gel (Bio-Rad) (30 µL, 18 well) and run for 1 hr at 140V in freshly diluted 10x Tris/Glycine/SDS running buffer (Bio-Rad, #1610732). Total protein using the stain-free gel imaging setting on the ChemiDoc MP was measured and then a semi-dry transfer using the high MW Midi setting (2.5A, 25V, 10 mins) onto methanol activated 0.2 µm PVDF membranes using cold 1x Trans-Blot Turbo Transfer Buffer with 20% ethanol (Bio-Rad, 10026938) was run. Blocking was done for 1h using AdvanBlock-Chemi Blocking Solution (R-03726-E10, Advansta) followed by a 15 min incubation in 1:20,000 streptavidin-HRP (Thermo Fisher, N100) in AdvanBlock-Chemi Blocking Solution (Advansta Inc). Streptavidin HRP solution was quickly removed, and the membrane was rinsed with DI water four times and washed three times with 1x PBST for at least 15 min. Finally, Clarity Western ECL Substrate (Bio-Rad, 1705060) was used for developing the membrane, and blots were imaged on a ChemiDoc MP (Bio-Rad).

#### Streptavidin pulldown and on-bead digestion

Precipitated proteins were pelleted by centrifugation at 4,000x g for 10 min at 4°C. Supernatant was aspirated and washed with 1mL of cold methanol:chloroform (4:1) mixture and then protein pellet was resuspended by water-bath sonication. This step was repeated once more. Finally, the proteins were pelleted at 4,000x g for 10 min at 4°C, the supernatant was aspirated and further air dried. The resulting dried protein pellets were solubilized in 10 µL of 10% SDS in DPBS, supplemented with 500 µL of freshly prepared 6M urea, and water bath sonicated until completely dissolved. Proteins were reduced by adding 50 µL using a freshly prepared, premixed solution of TCEP (25 µL, 200 mM in DPBS) and potassium carbonate (K_2_CO_3_, 25 µL, 600 mM in DPBS) for 45 min at 37°C with constant shaking. Free sulfhydryl groups were alkylated in the dark by adding 70 µL of freshly prepared iodoacetamide (IAA, 400 mM in PBS) for 30 min at room temperature. Following alkylation, samples were diluted with 130 µL of 10% SDS and 5.5 mL of PBS, then incubated with pre-washed 50% streptavidin agarose beads (100 µL of resin slurry per sample) for 2h at room temperature under end-over-end rotation. The beads were washed once with 5 mL of 0.2% SDS in PBS, twice with 5 mL of DPBS, once with 5 mL of Milli-Q water, by spinning down at 400x g for 5 min and supernatant was aspirated. Beads were transferred to a 2 mL protein lo-bind tube (Eppendorf) by making a slurry with 1mL of 100 mM TEAB, pH 8.5. Beads were spun down at 400x g for 5 min and supernatant was aspirated. Beads were resuspended in 100 µL of 100 mM TEAB with 1 mM CaCl_2_ and 1 µg of trypsin/Lys-c protease mixture. On-bead digestion was carried out overnight (∼16 h) at 37°C in the dark.

#### TMT-labeling and high pH-reverse phase fractionation

Following overnight digestion, beads were pelleted at 600x g for 5 min, and peptide-containing supernatant (∼100 µL) was carefully transferred to a new protein lo-binding tube. The beads were washed with 50 µL of freshly prepared 100 mM TEAB and pooled with the original supernatant, along with 60 µL Anhydrous acetonitrile (ACN). Peptides were labeled with 200 µg TMT isobaric tags (20 µg/µL in ACN, 10 µL added to samples) from a TMT10plex reagent set (Thermo Fisher, 90110) and incubated at RT for 1h with occasional vortexing. The labeling reaction was quenched by adding 8 µL 5% hydroxylamine in water for 15 min, after which the samples were acidified using 6 µL formic acid to stabilize the peptides. Labeled peptides evaporated to near-dryness in a speedVac centrifuge (Labconco Inc.) and stored at −80°C prior to fractionation. Peptides were reconstituted in 0.1% TFA and loaded onto a conditioned reverse phase spin column supplied in the Pierce™ High pH Reversed-Phase Peptide Fractionation Kit (Pierce, 84868), following manufacturer’s recommendations to condition the column. After loading, samples were spun down at 3,000x g for 1 min and the flow through was reloaded onto the column a second time and spun down. Once loaded, the column was washed three times with MS-grade water to remove salt and once with 5% ACN in 0.1% TEA buffer (supplied with the kit) to remove unreacted TMT reagent. Peptides were then eluted off the column in 16 fractions (300 µL per fraction) at increasing % ACN diluted in 0.1% TEA (7.5%, 10%, 12.5%, 15%, 17.5%, 20%, 22.5%, 25%, 27.5%, 30%, 32.5%, 35%, 40%, 45%, 50%, 60%). Fractions were then concatenated by combining the following pairs (1:9, 2:10, 3:11, 4:12, 5:13, 6:14, 7:15, 8:16), then the pooled fractions were frozen on dry ice then dried using a speedVac at 35°C.

LC-MS acquisition: A Dionex Ultimate Rapid Separation Liquid Chromatography system (Thermo Fisher) was used to load 10 µl of the reconstituted peptides (in 5% acetonitrile, 0.1% formic acid) onto a C18 trap column (Thermo Fisher) with a flow rate set at 5 µl/min for 10 min. Peptides were separated by reversed-phase chromatography on a 25-cm-long C18 analytical column (New Objective) packed in-house with BEH C18,130 Å, 1.7-um particle size (Waters) encapsulated in a column heater (MSWIL) with a temperature set at 60°C. Peptides were eluted by changing the mixture of mobile phase A (0.1% formic acid in water) and mobile phase B (0.1% formic acid in acetonitrile). The gradient program consisted of holding mobile phase B at 2% for the first 6 min, slowly ramped up to 35% over the next 124 min, followed by an increase to 85% over 5 min with a 5 min hold. The analytical column was re-equilibrated for 10 min prior to the next sample injection. The flow rate throughout the gradient was set to 0.3 µl/min. Eluted peptides were analyzed using an Orbitrap Eclipse Tribrid mass spectrometer (Thermo Fisher). The cycle time was set at top-speed for 1s with an MS1 mass scan range of 375 to 1,800 m/z and a mass resolution of 50,000. The most abundant precursor ions were fragmented with Higher Energy Collisional Dissociation (HCD) and with a collisional energy set to 38%. Dynamic exclusion was enabled for 60 s with a mass tolerance of ±10 ppm. The normalized AGC target was set to 250%. MS2 fragments were detected in the Orbitrap with a mass resolution of 50,000 with an injection time set to auto, and the MS2 mass scan range between 110 to 1800 m/z.

#### Mass spectrometry database search and analysis

Raw files were searched using MaxQuant 2.8.0.0 against a *Homo sapiens* UniProt database containing canonical sequences. Batch-specific TMT label correction factors were used, and reporter mass tolerance was set to 0.003 Da. Methionine oxidation, N-terminal acetylation, and asparagine/glutamine deamidation were allowed as variable modifications, and cysteine carbamidomethylation was allowed as a fixed modification. First search and main search peptide tolerance were both set to 20. Proteases were set to trypsin/P and LysC, with up to two missed cleavages allowed. Minimum peptide length was set to 6. PSM and protein FDR were both set to 0.01 with a minimum of 1 peptide, 1 razor+unique peptide, and 0 unique peptides. Protein group IDs were filtered to remove reverse sequences, contaminant sequences, and any proteins identified in < 8 samples. In the rare event of a zero value for a protein, the minimum non-zero value across all samples replaced the zero value. Differential abundance analysis was performed using the lmfit function from the limma package in R to fit a linear model to the log2 transformed abundance intensity of each protein group. The eBayes function was used to compute the moderate p-value and the topTable function to calculate adjusted p-value (using Benjamini-Hochberg multiple hypothesis correction) and FDR.

#### Transcript features

From the 6 h Emetine timepoint results, protein groups mapping to more than one gene symbol were expanded to one row per symbol. Emetine targets were defined as adjusted *P* < 0.05 and log_2_FC < −0.7 (*n* = 52); all other quantified proteins formed the background set (*n* = 5,195). Gene symbols were mapped to an Ensembl 96 human transcriptome annotation containing one transcript per gene, the isoform with the longest coding sequence, together with 5′UTR, CDS, and 3′UTR boundaries. Analysis was restricted to protein-coding genes with complete region boundaries (*n* = 51 targets, 5,131 background); 5′UTR, CDS, and 3′UTR sequences were extracted as substrings of the corresponding cDNA.

Region lengths were compared between sets using two-sided Wilcoxon rank-sum tests. Effect sizes are Cliff’s δ with 95% percentile confidence intervals from 2,000 bootstrap resamples; |δ| < 0.147 was taken as negligible (Romano et al., *Annual Meeting of the Florida Association of Institutional Research*; 2006).

#### Amino acid composition

Coding sequences were translated using the standard genetic code, and the percentage of each of the 20 standard amino acids was computed per protein. For each residue, the difference in mean percentage between targets and background and its 95% confidence interval were obtained from a two-sided Welch’s t-test; these are the values plotted. P values were adjusted across the 20 residues using the Benjamini–Hochberg procedure.

### Statistics and data availability

Data were analyzed using GraphPad Prism (v.11.1.0) unless otherwise indicated. Data are presented as mean ± SEM. Statistical analyses were performed using the Welch’s t-test, paired t-test, Wilcoxon test and one-way analysis of variance (ANOVA) with multiple comparison tests as stated. Blinding of treatment groups was performed for all imaging and subsequent analyses. Raw and processed scRNA-Seq and bulk RNA-Seq data have been deposited to GEO under accession number: GSE345453.

**Fig. S1:**
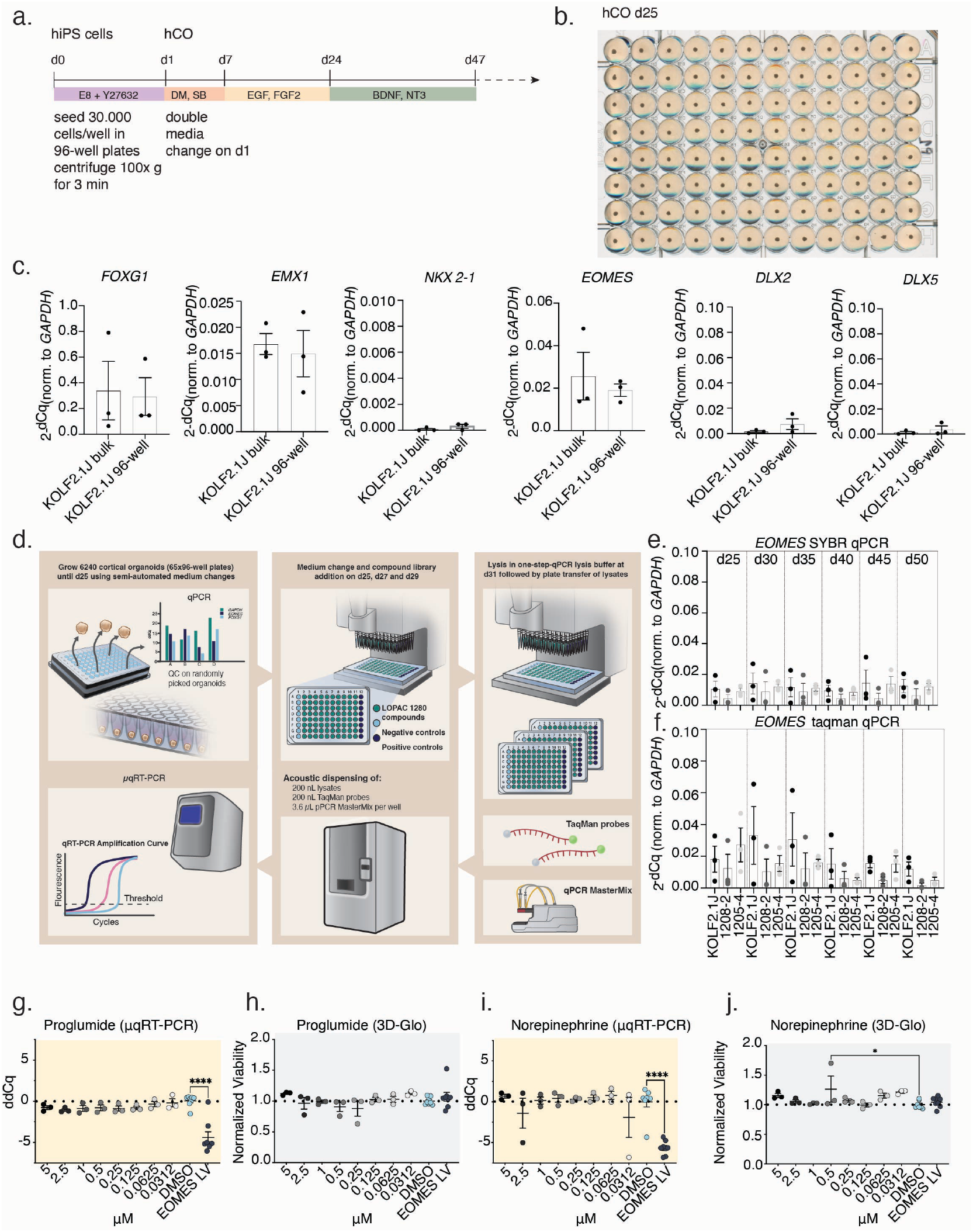
Development, optimization, and secondary validation of the 96-well hCO screening platform. **a)** Schematic detailing culture conditions for 96-well hCO. **b)** Representative image of a 96-well plate with hCO at 25 days of neural differentiation. **c)** qRT-PCR comparison of quality control markers for dorsal hCO patterning in 96-well or bulk grown hCO at 25 days. Expression of each marker was normalized to *GAPDH* and calculated as 2^-dCT^ (n=3 differentiations); all n.s. Two-tailed Welch’s t-test. **d)** Schematic describing the workflow of the LOPAC1280 small molecule screen using hCO. **e)** *EOMES* expression levels for hCO from day 25 to 50 *in vitro*, quantified by SYBR-based qRT-PCR (n=3 hiPS cell lines). Data are shown as mean ± SEM. **f)** *EOMES* expression levels for hCO from day 25 to 50 *in vitro*, quantified by Taqman-based qRT-PCR (n=3 hiPS cell lines). Data are shown as mean ± SEM. **g)** Dose response validation of induction of *EOMES* expression by Proglumide in the secondary screen. Each dot represents an individual well quantified by μqPCR and reported as ddCq relative to DMSO controls. *EOMES* overexpression is used as a positive control for the assay. Ordinary one-way ANOVA followed by Dunnett’s multiple comparisons test; F(9,30) = 9.699, P = <0.0001; EOMES LV ****P = <0.0001. Data are represented as mean ± SEM. **h)** Dose response assessment of hCO viability following Proglumide treatment in the secondary screen. Each dot represents an individual well quantified by 3D-Glo viability assay normalized to DMSO-treated controls. Ordinary one-way ANOVA followed by Dunnett’s multiple comparisons test; F(9,30) = 1.251, P = 0.3028. Data are represented as mean ± SEM. **i)** Dose response validation of reduction of *EOMES* expression by Norepinephrine in the secondary screen. Each dot represents an individual well quantified by μqPCR and reported as ddCq relative to DMSO controls. *EOMES* overexpression is used as a positive control for the assay. Ordinary one-way ANOVA followed by Dunnett’s multiple comparisons test; F(9,30) = 8.283, P = <0.0001; EOMES LV ****P = <0.0001. Data are represented as mean ± SEM. **j)** Dose response assessment of hCO viability following Norepinephrine treatment in the secondary screen. Each dot represents an individual well quantified by 3D-Glo viability assay normalized to DMSO-treated controls. Ordinary one-way ANOVA followed by Dunnett’s multiple comparisons test; F(9,29) = 2.200, P = 0.0522; for 0.5 μM *P = 0.0251. Data are represented as mean ± SEM.

**Fig. S2:**
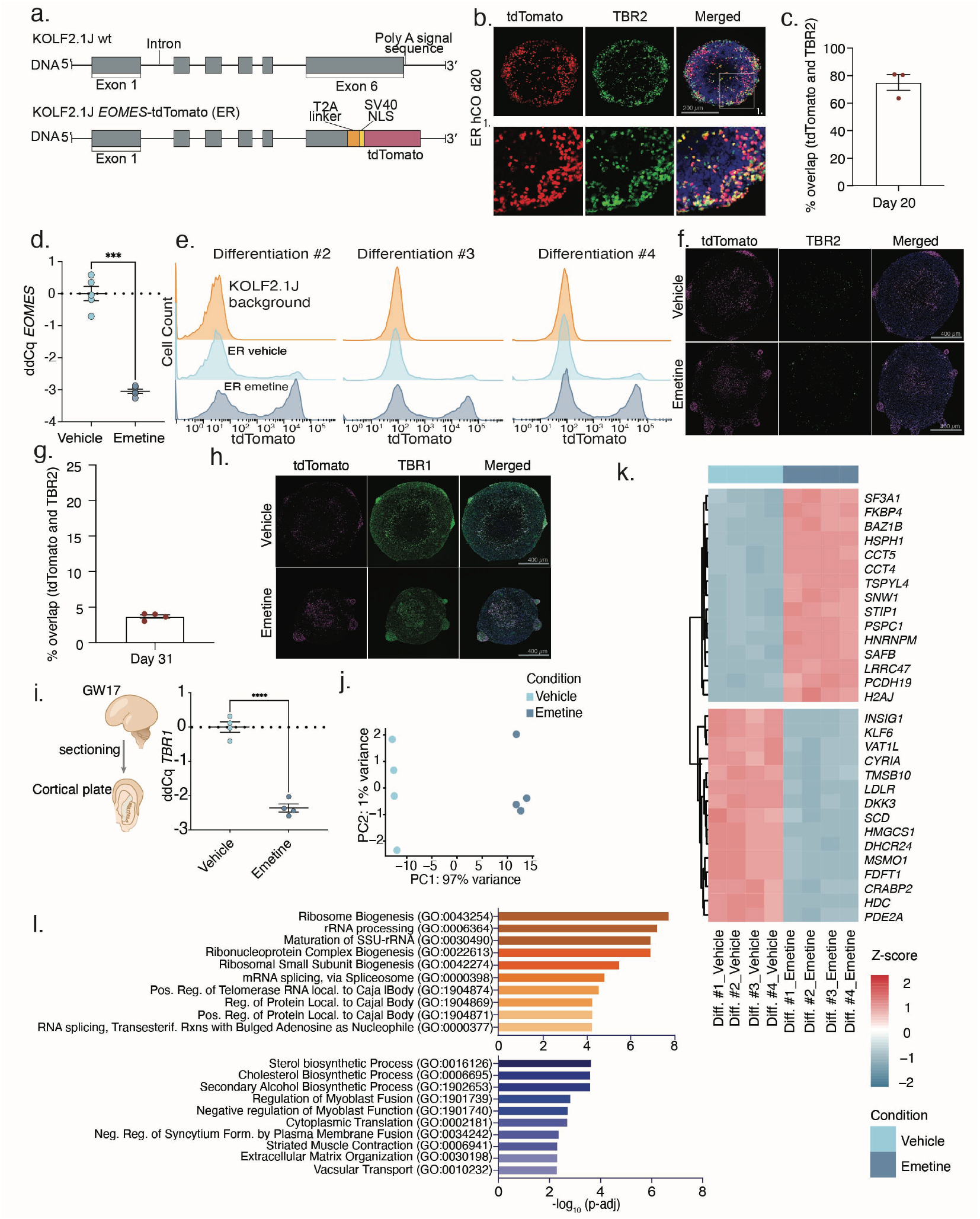
Characterization of the *EOMES* reporter line (ER) and ER hCO treated with Emetine. **a)** Generation of the *EOMES* reporter (ER) line in the KOLF2.1J hiPS cell background. **b)** Representative immunocytochemistry (TBR2) and fluorescence (tdTomato) images of ER hCO at 20 days of differentiation. tdTomato is shown in red, TBR2 in green and DAPI in blue. Scale bar = 200 μm. **c)** Quantification of percent overlap of tdTomato^+^ cells and TBR2 immunostained cells (n= 3 hCO). Data are represented as mean ± SEM. **d)** Validation of Emetine increase of *EOMES* expression in ER hCO. Each dot represents an independent differentiation (n=5 differentiations) experiment. *EOMES* expression was quantified by qRT-PCR and effect size is reported as ddCq relative to Vehicle-treated controls; Paired two-tailed t-test, ***P = 0.0002. Data are represented as mean ± SEM. **e)** FACS plots of ER hCO following Vehicle and Emetine treatment. Parental KOLF2.1J line derived hCO were used as a fluorescence background control. **f)** Representative immunocytochemistry (TBR2) and fluorescence (tdTomato) images of Vehicle and Emetine treated ER hCO at 31 days of differentiation. tdTomato is shown in magenta, TBR2 in green and DAPI in blue. Zoom out image of Fig. 2n. Scale bar = 400 μm. **g)** Quantification of percent overlap of tdTomato^+^ cells and TBR2 immunostained cells (n = 4 organoids). Data are represented as mean ± SEM. **h)** Representative immunocytochemistry (TBR1) and fluorescence (tdTomato) images of Vehicle and Emetine treated ER hCO at 31 days of differentiation. tdTomato is shown in magenta, TBR1 in green and DAPI in blue. Zoom out image of Fig. 2o. Scale bar = 400 μm. **i)** Schematic of *ex vivo* sliced human primary cortical cultures at GW17 and Emetine treatment of human primary cortical tissue. Expression of *TBR1* was quantified by qRT-PCR and effect size is reported as ddCq to Vehicle-treated controls. Each dot reports an independently cultured slice preparation (n=4 slices per condition); Welch’s two-tailed t-test, ****P = <0.0001. Data are represented as mean ± SEM. **j)** Principal component analysis (PCA) plot for dissociated FACS-isolated ER hCO following Vehicle or Emetine treatment (n= 4 independent differentiations for each condition). **k)** Heatmap of the top 15 differentially expressed genes ranked by differential expression p-value in FACS-sorted tdTomato^+^ cells from Vehicle and Emetine treated ER hCO at day 31 of neural differentiation. Expression values are shown as row-scaled Z-scores. **l)** Gene Ontology (GO) enrichment analysis of differentially expressed genes between Vehicle- and Emetine-treated FACS-sorted tdTomato^+^ cells. Genes with adjusted *P* ≤ 0.01 and |log2FC| > 1 were included in the analysis. Upper panel shows GO terms for upregulated genes and lower panel shows GO terms for downregulated genes.

**Figure S3:**
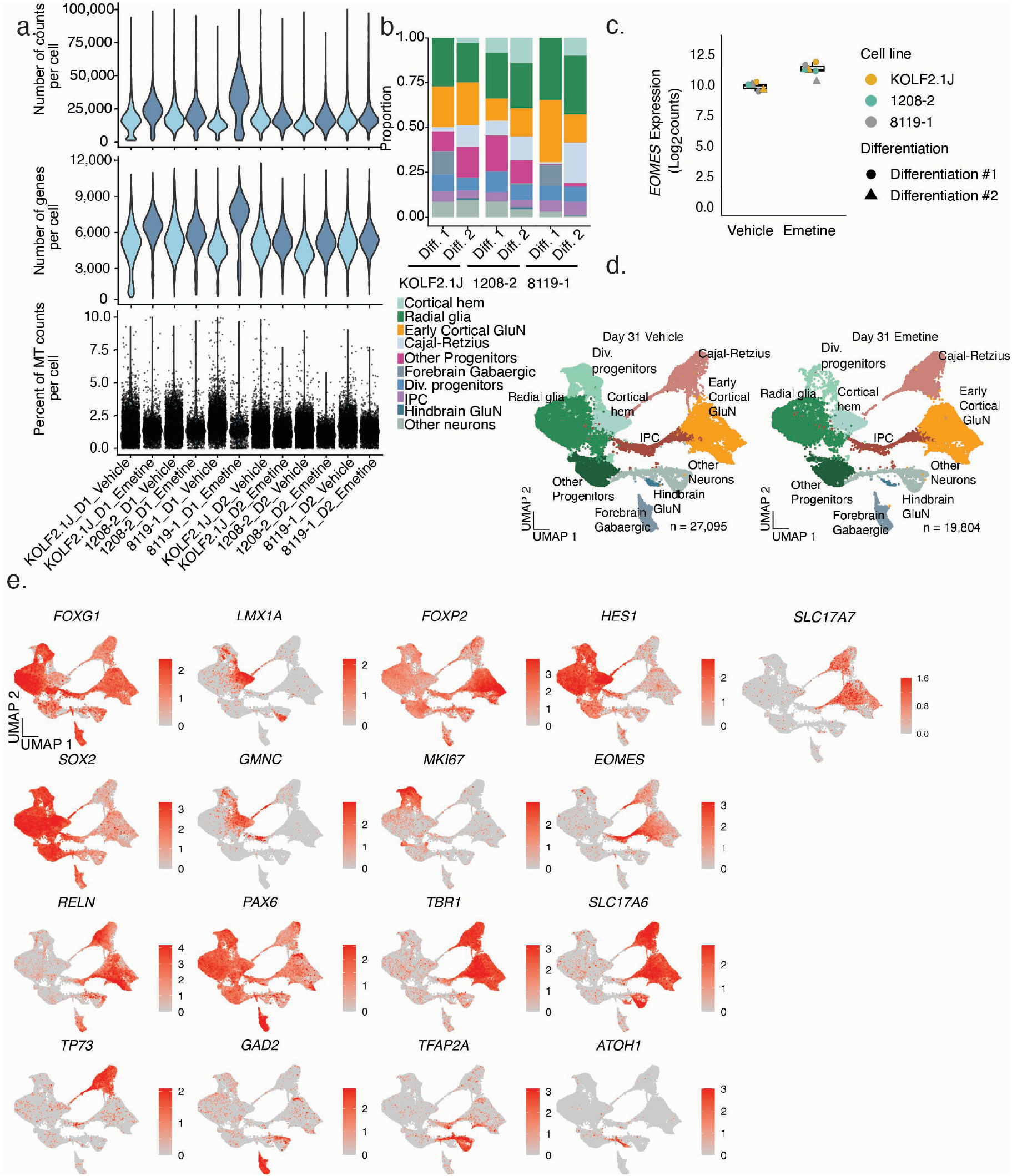
Single cell profiling of Emetine treated hCO at day 31 of neural differentiation. **a)** Quality control metrics for scRNA-Seq experiment at day 31 showing the distribution of the number of counts, number of genes, and mitochondrial (MT) gene fraction per cell in each sample. MT gene fraction plotted as boxplots. Horizontal line denotes median; lower and upper hinges correspond to the first and third quartiles; whiskers extend 1.5 times the interquartile range with outliers shown outside this range (n = 3 hiPS cell lines and two differentiations labeled as D1 and D2 for each condition). **b)** Bar plot displaying the proportion of annotated cell types in Vehicle and Emetine treated hCO at day 31 of differentiation. **c)** RNA expression (pseudobulk) normalized log_2_ counts for *EOMES* for Vehicle and Emetine treated hCO from scRNA-Seq profiling. **d)** UMAP visualization of single-cell gene expression of hCO at day 31 following Vehicle or Emetine treatment and cell type annotation (n = 27,095 cells for Vehicle treatment and n = 19,804 cells for Emetine treatment. Samples are from three hiPS cell lines and two differentiations for each condition). **e)** UMAP visualization of marker gene expression of Vehicle and Emetine treated hCO at day 31.

**Figure S4:**
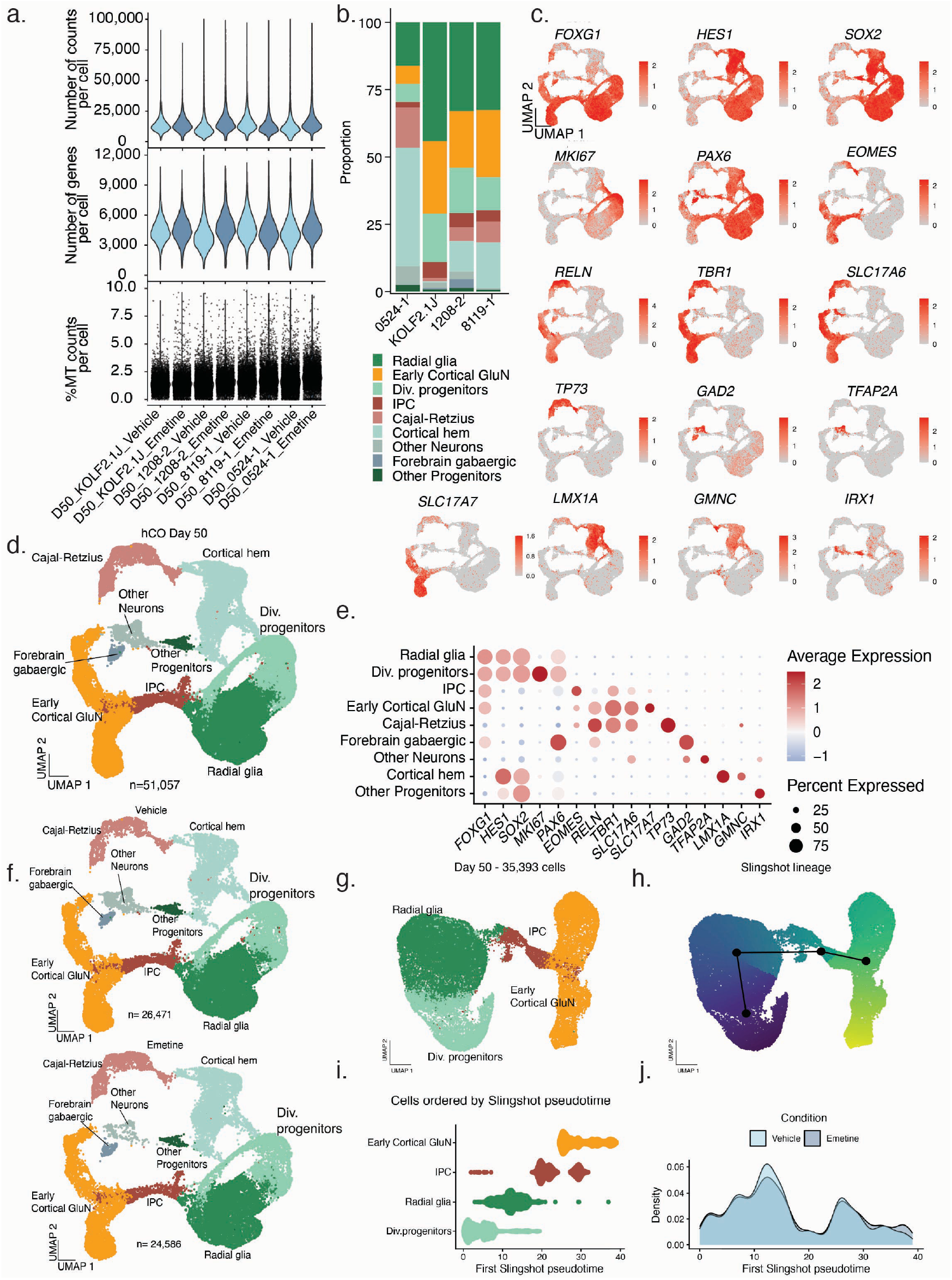
Single cell profiling of Emetine treated hCO at day 50 of neural differentiation to investigate longer term consequences of early exposure to Emetine. **a)** Quality control metrics for scRNA-Seq experiment at day 50 showing the distribution of the number of counts, number of genes, and mitochondrial (MT) gene fraction per cell in each sample. (n= 4 hiPS cell lines per condition). MT gene fraction plotted as boxplots (horizontal line denotes median; lower and upper hinges correspond to the first and third quartiles; whiskers extend 1.5 times the interquartile range with outliers shown outside this range). **b)** Bar plot displaying the proportion of annotated cell types in Vehicle and Emetine treated hCO at day 50 of differentiation. **c)** UMAP visualization of marker gene expression of Vehicle and Emetine treated hCO at day 50. **d)** UMAP visualization of single-cell gene expression of hCO at day 50 following Vehicle or Emetine treatment and cell type annotation (n = 51,057 cells from four hiPS cell lines). **e)** Dot plots of marker gene expression in Vehicle and Emetine treated hCO at day 50. The size of the circles represents the percentage of cells expressing the marker and color of the dots represent scaled gene expression values. **f)** UMAP visualization of single-cell gene expression of hCO at day 50 following Vehicle or Emetine treatment and cell type annotation (n = 26,471 cells for Vehicle treatment and n = 24,586 cells for Emetine treatment. Samples are from four hiPS cell lines for each condition). **g)** UMAP for subset of cells of dorsal forebrain lineage progression (n= 35,393 cells from 4 hiPS cell lines). **h)** Visualization of Slingshot lineage progression of dorsal forebrain cells at day 50 with associated pseudotime distributions along the lineage trajectory. **i)** Dorsal forebrain cell types ordered by pseudotime. **j)** Per-condition kernel density estimation of the pseudotime distributions at day 50. Permutation test P-value = 0.441.

**Figure S5:**
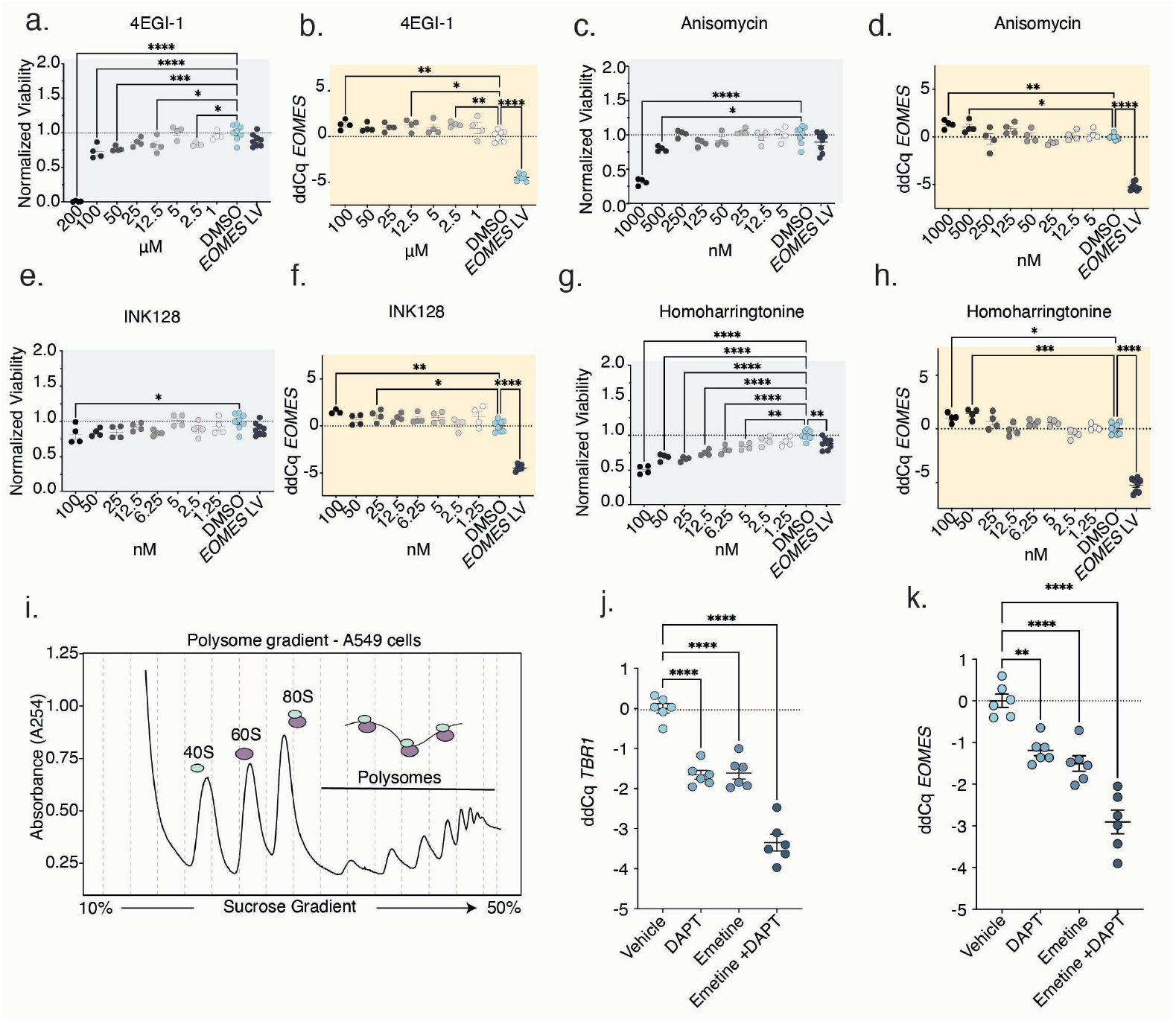
Characterization of translation inhibitors for *EOMES* expression in hCO and DAPT treatment. **a)** Dose response assessment of hCO viability following 4EGI-1 treatment in the translation inhibitor screen. Each dot represents an individual well quantified by 3D-Glo viability assay normalized to DMSO-treated controls. Ordinary one-way ANOVA followed by Dunnett’s multiple comparisons test; F(9,38) = 50.97, P = <0.0001; for 200 μM ****P= <0.0001, for 100 μM ****P=<0.0001, for 50μM ***P=0.0005, for 12.5 μM *P= 0.0121, for 2.5 μM *P= 0.0167. Data are represented as mean ± SEM. **b)** Dose response of *EOMES* expression by 4EGI-1 in the translation inhibitor screen. Each dot represents an individual well quantified by μqPCR and reported as ddCq relative to DMSO controls. *EOMES* overexpression is used as a positive control for the assay. Ordinary one-way ANOVA followed by Dunnett’s multiple comparisons test; F(8,35) = 69.19, P = <0.0001; for 100 μM **P= 0.0085, for 12.5 μM *P= 0.0116, for 2.5 μM **P= 0.0075, *EOMES* LV ****P = <0.0001. Data are represented as mean ± SEM. **c)** Dose response assessment of hCO viability following Anisomycin treatment in the translation inhibitor screen. Each dot represents an individual well quantified by 3D-Glo viability assay normalized to DMSO-treated controls. Ordinary one-way ANOVA followed by Dunnett’s multiple comparisons test; F(9,38) = 18.79, P = <0.0001; for 1000 nM ****P= <0.0001, for 500 nM *P=0.0132. Data are represented as mean ± SEM. **d)** Dose response of *EOMES* expression by Anisomycin in the translation inhibitor screen. Each dot represents an individual well quantified by μqPCR and reported as ddCq relative to DMSO controls. *EOMES* overexpression is used as a positive control for the assay. Ordinary one-way ANOVA followed by Dunnett’s multiple comparisons test; F(9,38) = 79.99, P = <0.0001; for 1000 nM **P= 0.0029, for 500 nM *P= 0.0297, *EOMES* LV ****P = <0.0001. Data are represented as mean ± SEM. **e)** Dose response assessment of hCO viability following INK128 treatment in the translation inhibitor screen. Each dot represents an individual well quantified by 3D-Glo viability assay normalized to DMSO-treated controls. Ordinary one-way ANOVA followed by Dunnett’s multiple comparisons test; F(9,38) = 2.504, P =0.0234; for 100 nM *P= 0.0275. Data are represented as mean ± SEM. **f)** Dose response of *EOMES* expression by INK128 in the translation inhibitor screen. Each dot represents an individual well quantified by μqPCR and reported as ddCq relative to DMSO controls. *EOMES* overexpression is used as a positive control for the assay. Ordinary one-way ANOVA followed by Dunnett’s multiple comparisons test; F(9,38) = 59.80, P = <0.0001; for 100 nM **P= 0.0038, for 25 nM *P= 0.0339, *EOMES* LV ****P = <0.0001. Data are represented as mean ± SEM. **g)** Dose response assessment of hCO viability following Homoharringtonine treatment in the translation inhibitor screen. Each dot represents an individual well quantified by 3D-Glo viability assay normalized to DMSO-treated controls. Ordinary one-way ANOVA followed by Dunnett’s multiple comparisons test; F(9,38) = 27.60, P = <0.0001; for 100 nM ****P= <0.0001, for 50 nM ****P= <0.0001, for 25 nM ****P= <0.0001, for 12.5 nM ****P= <0.0001, for 6.25 nM ****P= <0.0001, for 5 nM **P= 0.0013, for *EOMES* LV **P =0.0039. Data are represented as mean ± SEM. **h)** Dose response of *EOMES* expression by Homoharringtonine in the translation inhibitor screen. Each dot represents an individual well quantified by μqPCR and reported as ddCq relative to DMSO controls. *EOMES* overexpression is used as a positive control for the assay. Ordinary one-way ANOVA followed by Dunnett’s multiple comparisons test; F(9,38) = 92.32, P = <0.0001; for 100 nM *P= 0.0189, for 50 nM ***P= 0.0008, *EOMES* LV ****P = <0.0001. Data are represented as mean ± SEM. **i)** Representative polysome profile of A549 cell lysate as positive control. Lysates were separated by ultracentrifugation through a 10–50% sucrose gradient, and absorbance was measured at 254 nm. Peaks corresponding to the 40S and 60S ribosomal subunits, 80S monosomes, and polysomes are indicated. **j)** *TBR1* expression following Emetine and DAPT treatment of hCO. Each dot represents an independent differentiation (n=3 differentiations and two replicates each). TBR1 expression was quantified by qRT-PCR and effect size is reported as ddCq relative to Vehicle-treated controls. Ordinary one-way ANOVA followed by Dunnett’s multiple comparisons test; F(3,20) = 79.62, P = <0.0001; DAPT ****P =<0.0001, Emetine ****P = <0.0001, Emetine + DAPT ****P = <0.0001. Data are represented as mean ± SEM. **k)** *EOMES* expression following Emetine and DAPT treatment of hCO. Each dot represents an independent differentiation (n=3 differentiations and two replicates each). EOMES expression was quantified by qRT-PCR and effect size is reported as ddCq relative to Vehicle-treated controls. Ordinary one-way ANOVA followed by Dunnett’s multiple comparisons test; F(3,20) = 36.67, P = <0.0001; DAPT **P = 0.0011, Emetine ****P = <0.0001, Emetine + DAPT ****P = <0.0001. Data are represented as mean ± SEM.

**Fig. S6:**
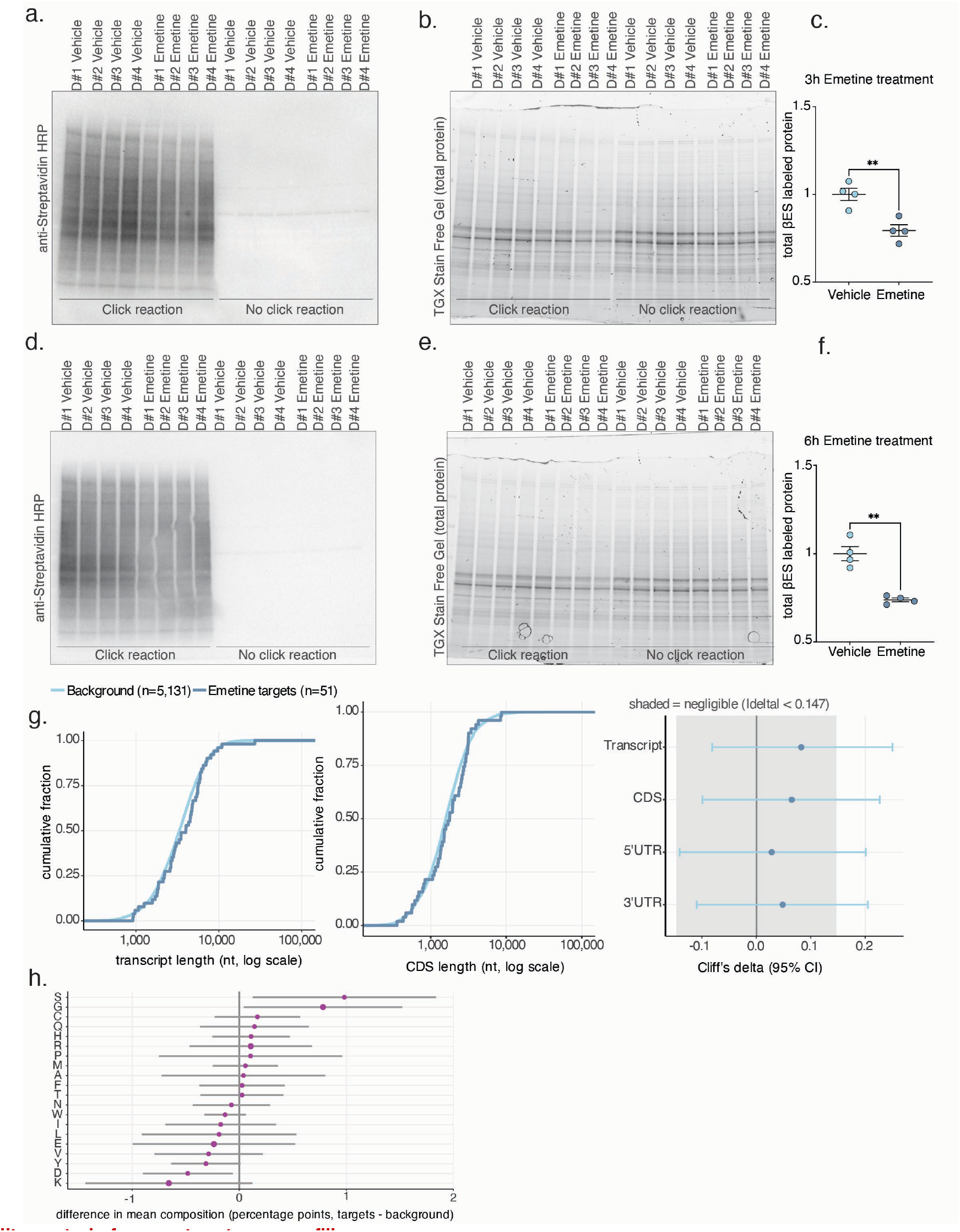
Quality control of nascent proteome profiling. **a)** Immunoblot of Streptavidin-HRP on all 3h samples from nascent proteome profiling. **b)** Stain-free gel image for depicting total protein amounts. **c)** Quantification of βES labeled nascent peptides relative to total protein after 3 hours of Emetine treatment (n=4 differentiations), Welch’s two-tailed t-test P= 0.0050. **d)** Immunoblot of Streptavidin-HRP on all 6h samples from nascent proteome profiling. **e)** Stain-free gel image for depicting total protein amounts. **f)** Quantification of βES labeled nascent peptides relative to total protein after 6 hours of Emetine treatment (n=4 differentiations), Welch’s two-tailed t-test P= 0.005. **g)** Emetine targets (adjusted P < 0.05, log_2_FC < −0.7; n = 51) were compared with all other quantified proteins (n = 5,131). (Left, Middle) Empirical cumulative distributions of transcript length (Left) and CDS length (Middle). Median transcript length was 4,004 nt in targets versus 3,332 nt in background (Wilcoxon P = 0.31); median CDS length was 1,773 versus 1,584 nt (P = 0.42). (Right) Cliff’s δ with 95% bootstrap confidence intervals for transcript, CDS, 5′UTR, and 3′UTR length; shading marks the negligible range (|δ| < 0.147). All point estimates fell within the negligible range and every interval included zero, with upper bounds reaching δ = 0.20–0.25. **h)** Difference in mean amino acid composition (percentage points, targets minus background; n = 51 and 5,131) for all 20 residues, with 95% confidence intervals with Benjamini–Hochberg correction across the 20 residues (minimum q = 0.26). All confidence intervals excluded differences larger than ±1.9 percentage points.

**Movie S1**: Representative z-projected live confocal video of Vehicle and Emetine treated ER hCO from day 4 to day 6 of Emetine treatment. Scale bar = 200 μm.

